# Integrated analysis of high throughput transcriptomic data revealed specific gene expression signature of cardiomyocytes

**DOI:** 10.1101/2020.06.14.151399

**Authors:** Mohammad Reza Omrani, Erfan Sharifi, Niusha Khazaei, Sahel Jahangiri Esfahani, Nicholas W. Kieran, Hadi Hashemi, Abdulshakour Mohammadnia, Moein Yaqubi

**Affiliations:** National Institute of Genetics Engineering and Biotechnology (NIGEB), Tehran, Iran; Department of Biology, Université de Sherbrooke, Sherbrooke, QC, Canada; Department of Human Genetics, Faculty of Medicine, McGill University, Montreal, Quebec, Canada; Neuroimmunology unit, Montreal Neurological Institute, McGill University Montreal, QC; Shiraz University of Technology, Department of Electrical and Electronic Engineering, Shiraz, Fars, Iran; Integrated Program at Neuroscience, Neuroimmunology unit, Montreal Neurological Institute, McGill University Montreal, QC, Canada

**Author notes:** these authors contributed equally to this work (Mohammad Reza Omrani, Erfan Sharifi). **Corresponding authors:** Abdulshakour Mohammadnia Research Institute of the MUHC, 1001 Decarie Boulevard, Montreal, QC H4A 3J1, Canada, Moein Yaqubi 3801 Rue Université, Montréal, QC H3A 2B4, Canada.

**Keywords:** cardiomyocytes, differentiation, heterogeneity, High throughput sequencing, transcriptome

## Abstract

Acquiring a specific transcriptomic signature of the human and mouse cardiomyocyte (CM) will greatly increase our understanding of their biology and associated diseases that remain the most deadly across the world. In this study, using comprehensive transcriptomic mining of 91 cell types over 877 samples from bulk RNA-sequencing, single cell RNA-sequencing, and microarray techniques, we describe a unique 118-gene signature of human and mouse primary CMs. Once we had access to this CM-specific gene signature, we investigated the spatial heterogeneity of CMs throughout the heart tissue. Moreover, we compared the CM-specific gene signature to that of CMs derived from 10 differentiation protocols, and we identified the protocols that generate cells most similar to primary CMs. Finally, we looked at the specific differences between primary and differentiated CMs and found that differentiated cells underexpress genes related to CM development and maturity. The differentiated cells conversely overexpressed cell cycle-related genes, resulting in the progenitor features that remain in differentiated CMs compared to primary adult CMs. The presence of histone post translational modification H3K27ac from ChIP sequencing data sets were used to confirm transcriptomic findings. To the best of our knowledge, this is the most comprehensive study to date that unravels the unique transcriptomic signature of primary and differentiated CMs. This study provides important insights into our understanding of CM biology and the molecular mechanisms that make them such a unique cell type. Moreover, the specific transcriptomic signature of CMs could be used in developmental studies, stem cell therapy, regenerative medicine, and drug screening assays.

## Introduction

Cardiomyocytes (CMs) constitute the majority of heart by mass and have an essential role in tissue contraction and overall heart function.^1^ CM damage and dysfunction is the primary cause of heart-related diseases. Despite decades of research related to drug development, regenerative medicine, and cell therapy for heart-related disorders, heart disease remains the global leading cause of death and has seen minimal progress in treatment strategies.^2,3^ Gene expression quantification of CMs is a crucial step to understand their contribution to disease development, which could lead to new therapies.^4^ This transcriptomic profile of primary CMs could be used as a reference to guide differentiation protocols of induced CMs. More humanistic CMs would then be invaluable for research, drug screening studies, and regenerative medicine. In spite of rich literature exploring the potential uses of differentiated CMs, there is an unmet need to define the unique transcriptomic signature of these cells that would open up their full potential.

Bulk RNA-sequencing is the most commonly used technique in transcriptomic research. While it provides detailed information on the entire tissue under investigation (heart tissue in this case), single cell RNA-sequencing is required to isolate individual cells and look exclusively at CMs. In one recent paper, Wang and colleagues identified different subtypes of human CMs in the left atrium and ventricle.^5^ Moreover, spatio-temporal heterogeneity of mouse CMs has been studied by DeLaughter et al. (2016), in which they discovered that the CM subtype depends on the stage of heart development and the location of the heart tissue.^6^ Other single cell RNA-seq studies have further noted the heterogeneity of cardiac muscle cells, with a gradient expression of CM marker genes like ACTC1 and MYH6.^7^ Moreover, single cell studies have revealed the diverse cellulome of heart tissue through identifying major cell types and their gene expression heterogeneities in the heart.^8^ Altogether, despite improvements in our understanding of the transcriptional profile of the heart and CMs, larger studies that combine hundreds of datasets are required to confidently define the specific gene expression profile of CMs compared to other cells in the body.

Heart transplant is currently the treatment of choice for end-stage heart failure; however, it is an unsustainable and rare solution given the scarcity of donors that limits this treatment to a small portion of patients.^9^ To overcome such obstacles, other strategies have been proposed, including stimulating the proliferation of existing CMs, activation of endogenous cardiac progenitor cells, and differentiation of pluripotent stem cells or direct reprogramming of somatic cells towards CMs.^10–12^ Among these techniques, generation of CMs through stem cell differentiation and somatic cell direct reprogramming have earned the most attention over the past decade. For example, van den Berg and colleagues proposed a monolayer differentiation technique through which pluripotent stem cells can be converted to CMs.^13^ Similarly, it has been indicated that mouse fibroblasts can be converted directly into cardiac muscle cells using three transcription factors, Gata4, Tbx5 and Mef2c.^14^ To assess the maturity of differentiated CMs, the transcriptomic profile is supplemented by the structure and function of the cytoskeleton and contractile apparatus, metabolic status, and electrophysiological properties.^15–18^ These studies also show that, despite significant progress in the generation of more mature CMs, they still fail to functionally resemble their adult *in vivo* counterparts. Therefore, determining the transcriptomic differences between *in vivo* and differentiated CMs through comprehensive transcriptomic analysis will allow for more targeted differentiation techniques to create CMs more useful for research and eventually regenerative medicine.^19^

In this study, we have conducted comprehensive transcriptomic analysis of 877 gene expression samples complemented with epigenetic data sets to define the specific transcriptomic profile of CMs. In addition, we assessed spatial gene expression heterogeneity of human CM populations. Furthermore, we used our source of data to precisely compare several differentiation protocols and show the similarity of differentiated cells with their *in vivo* counterparts. Finally, through assessing 15 different differentiation studies, we found common transcriptomic features of all pluripotent stem cell-derived CMs that were significantly different from mature adult human CMs. Our transcriptomic finding well correlated with H3K27ac peaks derived from ChIP-seq data sets. Our results, in addition to identifying the fundemental transcriptomic features of cardiac muscle cells, could be crucial in future translational studies, like generating mature CMs for cell based therapy and tissue engineering studies.

## Results

To characterize the specific transcriptomic profile of cardiomyocytes (CMs), we analyzed human and mouse transcriptomic data of 91 somatic cell types and subtypes, as well as multiple types of stem cells (65 human and 26 mice cell types). We categorized cells based on the cell type, tissue and organ (location), and developmental stages (embryonic/fetal, pediatric/adolescent, and young/adult). The transcriptomic data came from bulk RNA-seq, microarray, and single cell RNA-seq experiments. By analyzing the compiled data sets, we identified a unique profile of genes that was differentially expressed in CMs compared to all other cell types. We analyzed this specific profile of genes to investigate their function and heterogeneity in CMs. In addition, we assessed the success of CM differentiation from human pluripotent stem cells in a number of differentiation protocols to find the most efficient protocols in generating CMs similar to their adult human counterparts. Finally, we compared differentiated CMs to primary adult CMs to explore transcriptomic differences that may hinder the application of differentiated CMs in research and regenerative medicine.

### Comprehensive transcriptome mining reveals cardiomyocyte-specific gene expression landscape

To gain insight into the transcriptomic profile that make CMs a distinct cell type with highly specialized functions, we compiled data from 180 studies that examine 91 different cell types or subtypes from 877 individual RNA-seq and microarray experiments (Fig. 1A, Fig. S1A). All samples were curated to remove outliers. Expression of canonical CM marker genes were checked in the compiled CM dataset and a consistent expression of the marker genes was seen in them which implies the high quality of the gathered CM samples (Table S1 and Fig. S2A). Comparing RNA-seq and microarray data sets of human and mouse CMs to all other somatic and stem cell types revealed 137 genes with significantly higher expression levels in both mouse and human CMs compared to any other cell types. Because the data in Fig 1A is generated from bulk RNA-seq and microarray techniques, which are unable to capture transcriptomic information of individual cells, we checked the validity of the identified genes at the single cell level. For single cell gene expression analysis, we used the Tabula Muris Consortium, which contains mice single cell RNA-seq data.^20^ Cross-referencing our 137 upregulated genes from bulk RNA-seq and microarray data to the Tabula Muris Consortium identified 118 of the genes to also be significantly upregulated in CMs at the single cell level. Jaccard similarity correlation analysis confirms that these 118 genes make primary CM to cluster together and apart from all other human RNAseq datasets (Fig. S2B). Due to the higher expression of these genes in CMs, these 118 genes functioned as identity markers of CMs compared to other cells (Fig. 1B and Table S2). Across all cell types, the expression of housekeeping genes remained relatively constant, with the exception of ACTB that had lower expression in CMs compared to somatic cells (Fig. 1B). This observation suggests that the significantly upregulated genes in this study are highly specific for CMs and are not artefacts generated from the acquisition of transcriptomic data. As expected, gene ontology analysis and human phenotype ontology revealed the heavy contribution of these genes to muscle development and in the normal function of the heart, as their dysregulation results in heart-related abnormalities like myopathy (Fig. 1C and 1D).

**Figure 1.**
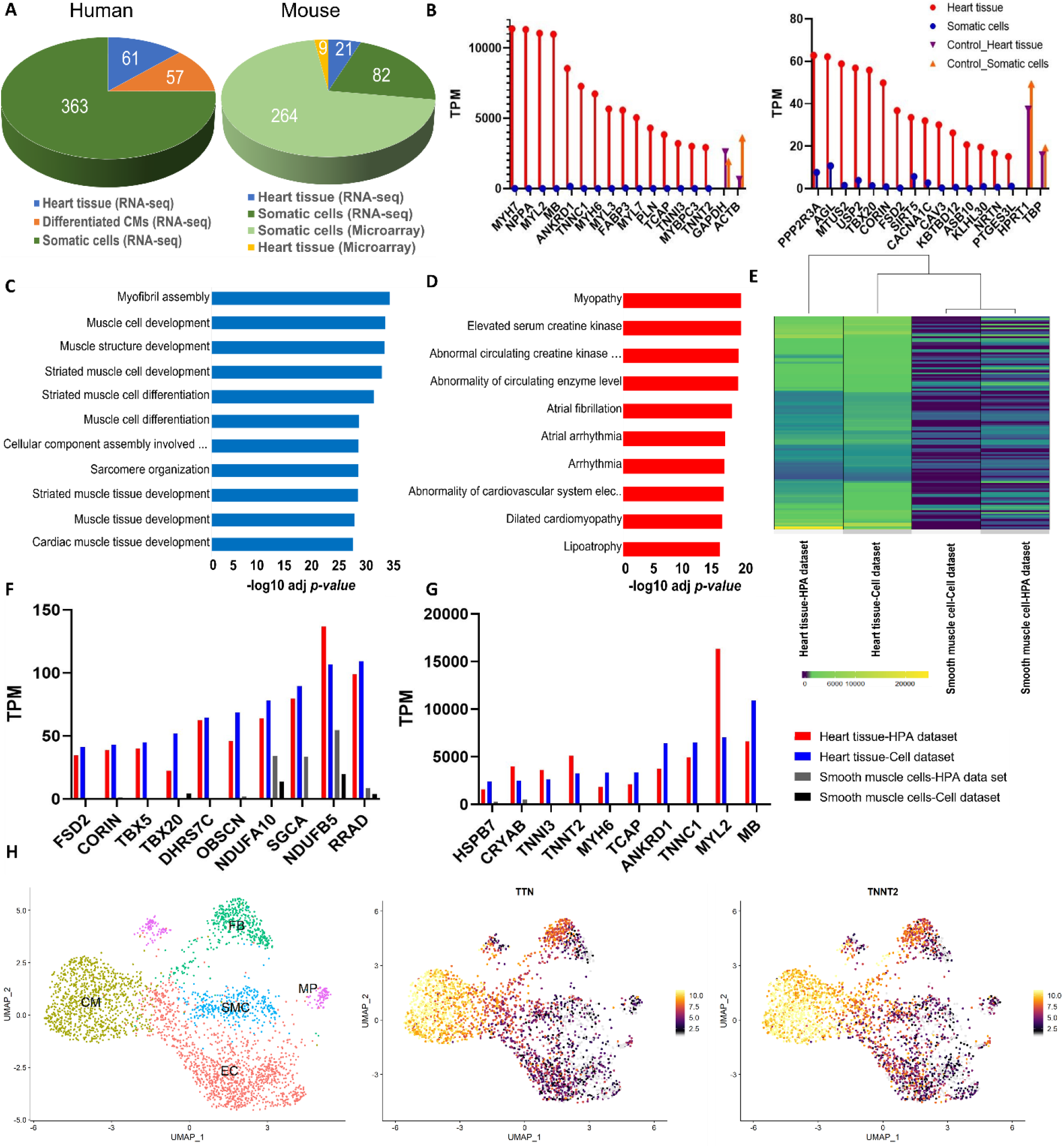
A) The distribution of transcriptomic analysis type (RNA-seq or microarray) are described for both human and mouse samples. B) Bar graphs quantifying the highest and lowest TPM values of cardiomyocyte (CM)-specific genes. Left panel: expression of the 15 genes with the highest TPM values in CMs. Right panel: expression of the 15 genes with the lowest TPM values in CMs. Expression of GAPDH, ACTB, HPRT1 and TBP were used as reference genes. C & D) Gene ontology and human phenotype ontology analysis of CM-specific genes were ordered based on their −log10 adjusted *p-value*. E) A heatmap showing the expression levels of the118 CM-specific genes in the heart tissue and smooth muscle cells used in this study, as well as in the heart tissue and smooth muscle cells obtained from the human protein atlas (HPA) data set. Higher expression of the genes is represented with yellow, while lower expression of genes is represented with dark blue. F & G) Quantification of the expression levels of genes in the heart tissue and smooth muscle cell generated from our individually collected samples and the HPA data set. Lowly expressed genes are represented in (F) and highly expressed genes are expressed in (G). H) The UMAP of reanalyzed single cell RNA-seq data set of Wang et al., study of 3,168 CMs and other non-CM cells. Among the non-CM cells were fibroblasts (FB), macrophages (MP), endothelial cells (EC), and smooth muscle cells (SMC). Expression of TTN and TNNT2 are shown as examples of CM-specific genes. The expression of genes is indicated by a color gradient in which yellow show high and black shows low expression.

To test and validate the reproducibility of our findings, we recruited publicly available important transcriptomic source for human tissue and organs, the Human Protein Atlas (HPA) dataset.^21^ Although this source contains whole tissue and organs data (not individual cells), it remains valuable resource to which we can compare our human and mouse transcriptomic data. We extracted the expression level of 118 CM-specific gene list in the HPA data set which contains expression data of 42 different tissues and made an expression matrix which contain 4956 data points (each data point correspond to one single expression value in the matrix). Interestingly, in 98.06% data points (4860 out of 4956) expression of these 118 genes were higher in heart tissue compared to all other tissues which indicates the reliability of our findings. The most similar tissue to the heart tissue in this matrix is skeletal muscle cell. Even though we did not have the skeletal muscle cell information in our study, if we look at the expression of 118 cells individually between the heart and skeletal muscle tissues in the HPA database, we see 69 genes have higher expression in the heart compared to skeletal muscle cells. However, principle component analysis (PCA) analysis using expression value of 118 genes shows a clear separation between CMs and skeletal muscle cells (Fig. S1B). Therefore, we can confidently consider this expression signature as CM specific profile. So far, we considered expression of 118 CM-specific genes which is provided by HPA (intra-database analysis). At the next step we wanted to directly compare the expression values that we get in our analysis with the HPA information (Inter-database analysis). To this end, we chose smooth muscle cells and tissue as an example comparator which is a cell type and tissue which that is present in both datasets. The heart tissue and smooth muscle cell information of HPA were directly compared to heart tissue and smooth muscle cell information of our study respectively. While there was a high correlation between comparisons at tissue and cell level in which both data sets showed similar expression pattern (Fig. 1E), individual expression values were higher in CM compared to the smooth muscle cell (Fig. 1F and G). In conclusion, the high correlation between CMs and heart tissue from the HPA dataset, as well as and low correlation with smooth muscle HPA data, confirmed the reliability of our data to assess the unique gene expression signature of heart cardiac muscle cells.

To validate the specificity of the identified CM-specific genes in human heart tissue, we reanalyzed the available single cell RNA-seq data set from a Wang et al. (2020), in which they isolated and sequenced the heart tissue that included CMs and non-CM cells. Following the quality control and removing unwanted cells we ended up having 3168 cells including CMs (32%), fibroblasts (13%), smooth muscle cells (11%), endothelial cells (39%), and macrophages (5%) in the human heart tissue consistent with the findings of their original study. We checked the previously identified 118 CM genes in this single cell dataset, and all detected CM genes were again more highly expressed in the CMs than any other heart tissue cell types (Fig. 1H and Fig. S2C). Due to the low sensitivity of single cell RNA-seq, the expression of some genes could not be identified (Fig. S3). Overall, we analyzed an extensive array of transcriptomic data to confidently define a CM-specific gene expression profile and we compared this list to CM single-cell sequencing data to confirm the validity of the data.

In addition to measuring the mRNA expression of genes, identifying the level of histone post-translational modifications provides important insight into the accessibility of the gene for transcription. Post-translational modifications of histone tails, specifically acetylation at lysine 27 of histone H3 (H3K27ac), is tightly coupled and correlated with gene expression activation. In this regard, we used ChIP-seq data from human epigenome roadmap dataset to identify epigenetic signature of our identified genes and correlate these epigenetic signatures with their expression level.^22^ We observed enriched H3K27ac marks on the promoters of CM-specific genes, as an example we chose top 15 most highly expressed CM-specific genes to visualize the presence of H3K27ac peaks (Fig. 1B, first panel). The majority of the analyzed genes had a high correlation between increased expression and the presence of H3K27ac in their promoter regions (Fig. 2A). We also looked at the presence of H3K27ac on CM-specific genes in 14 other cell types and found that the acetylation peaks were specific to cells of the heart tissue (Fig. 2A). To control the validity of our analysis across different ChIP-seq experiments, we examined the H3K27ac marks of genes known to be upregulated in other cell types, including pluripotent specific genes (POU5F1/OCT4), a hepatocyte marker (ALB), and a smooth muscle marker (MYH11). The H3K27ac peaks were indeed present in the promoters of these genes in the correct cellular context and were not present in heart and other cell/tissue types (Fig. 2A, S4A). In addition, we used H3K27ac ChIP-seq experiment from left ventricle of adult female downloaded from ENCODE project to confirm identified peaks using human epigenome roadmap data (Fig. S4B). In summary, analysis of H3K27ac marks revealed a high level of specificity and a strong correlation between H3K27ac marks and CM-specific gene expression of heart cells.

**Figure 2.**
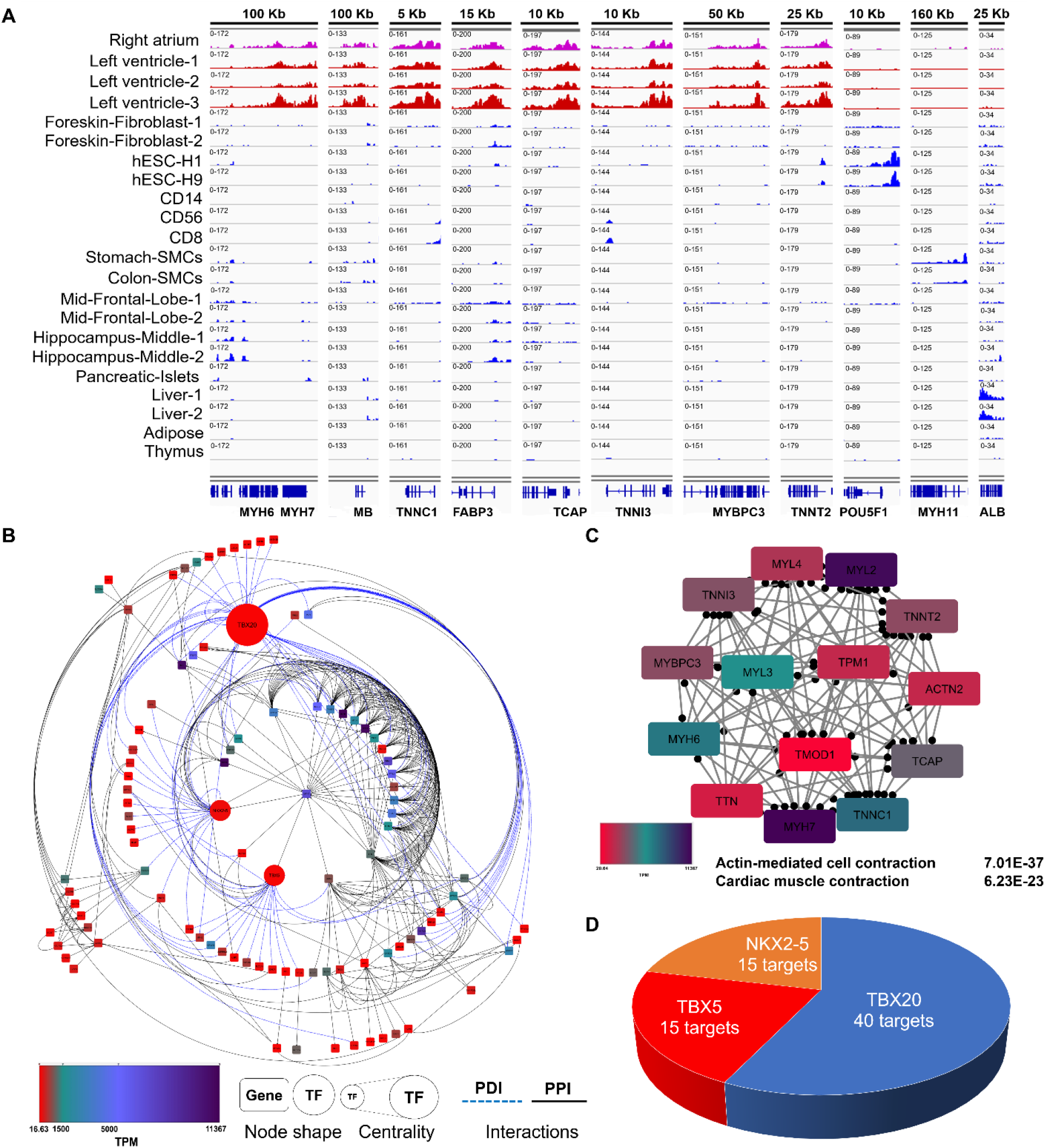
A) H3K27ac peaks on the promotor region of the genes across different cell and tissue types. B) Gene regulatory network of CM-specific genes using protein-protein (PPI) and protein-DNA (PDI) interaction data. 97 out of the 118 CM-specific genes are the nodes of the network and are connected by 311 interactions. Genes are shown as rectangles and TFs as circles. Red and dark blue colors show the genes or TFs with the lowest and highest TMP respectively. Also, solid lines denote PPIs while dashed lines denote PDIs. C) The most significant protein complex with 14 highly expressed genes is involved in actin-mediated cell contraction. Red and blue color indicate low and high expression of genes, respectively. D) Pie chart of the most important TFs. TBX20 with 40 targets and TBX5 and NKX2-5 each with 15 targets play crucial roles as regulators of downstream 118 genes.

To better understand the interaction of these specific genes and their role in establishing major CM functions, we constructed an integrated gene regulatory network using protein-DNA and protein-protein interaction data of the genes (Fig. 2B). The constructed network contains 97 of the CM-specific genes which have been connected by 311 protein-protein and protein-DNA interactions (Fig. 2B). Protein-protein interaction analysis revealed four significant complexes, one of which has 14 highly expressed CM genes that are involved in actin-mediated cell contraction (Fig. 2C, Fig. S5A, S5B, and S5C). Furthermore, regulatory network analysis revealed that three transcription factors in the list of 118 genes, TBX20, NKX2-5 and TBX5, regulate the expression of most downstream genes in the network. TBX20 alone regulates 40 genes in the network, while NKX2-5 and TBX5 each target 15 genes (Fig. 2D). In summary, by observing protein-protein and protein-DNA interactions, we have identified that many CM-specific genes are involved in cardiac muscle function and additionally, our results revealed that TBX20 is specific master regulator of this specific profile.

### Spatial heterogeneity of cardiomyocyte-specific transcriptome

Single cell RNA sequencing technology provides an opportunity to explore heterogeneity within a given cell type or lineage. Therefore, we investigated the spatial heterogeneity of the identified genes in the CMs of heart tissue. To address this aim, we used the Wang et al. (2020) study and analyzed single cell data of 12 normal heart samples (five samples from the left ventricle and seven samples from the left atrium).^5^ After removing unwanted and low-quality cells, we kept 4881 cells. Unsupervised clustering of the cells resulted in identification of 13 different clusters: six clusters for the left ventricle (LV), six clusters for the left atrium (LA), and one atrioventricular (AV) cluster based on ventricle and atrium markers reported in a Wang et al., (2020) study (Fig. 3A). Most of the CM-specific genes that we identified in the previous analysis (118 genes), like COX7A1, did not show a significant difference between LV and LA clusters (Fig. 3A). 19 genes did show significant differential expression between LV and LA (Fig. 3A). Among these 19 genes, MYH7, MYL2, MYL3, and TNNI3 were enriched in the LV, while 15 genes were enriched in the LA, including NPPA, NPPB, TRDN, CMYA5 and TTN (Fig. 3A, 3B, Fig. S6A and S6B). We looked into the H3K27ac pattern of these genes to see if we could find the same pattern of heterogeneity at the epigenetic level. Interestingly, we found that multiple genes had different H3K27ac patterns in the atrium compared to ventricle (Fig. 3C). For example, we found that the NPPA, MYL4 and MYL7 promoters had higher H3K27ac peaks in the right atrium compared to the left and right ventricles (Fig. 3C and Fig. S4B). We identified ventricle-specific genes MYL2 and MYL3 that had higher H3K27ac marks in the ventricles compared to the atrium (Fig. 3C, S4A). LA specific genes are mainly involved in the muscle cell function and muscle cell development (Fig. 3D). In addition to highlighting heterogeneity between LA and LV for CM-specific genes, we interestingly discovered that CM-specific genes are differentially expressed between different clusters of both the ventricle and atrium, with a continuum expression profile of many genes observed (Fig. 3E and 3F). We found 73 genes differentially expressed in LA1 versus other LA clusters, and 53 genes show significant variation in LV1 compared to other LV clusters. CMYA5 and MB are represented as examples of heterogeneity of the genes in the ventricle and atrium clusters respectively (Fig. 3 G and H). This observation may imply that the atrium has more dispersed gene expression than the ventricle. Altogether, our analysis showed that there is a spatial heterogeneity for CM-specific genes between the atrium and ventricle and, moreover, there is a gradient expression of these genes within each heart chamber.

**Figure 3.**
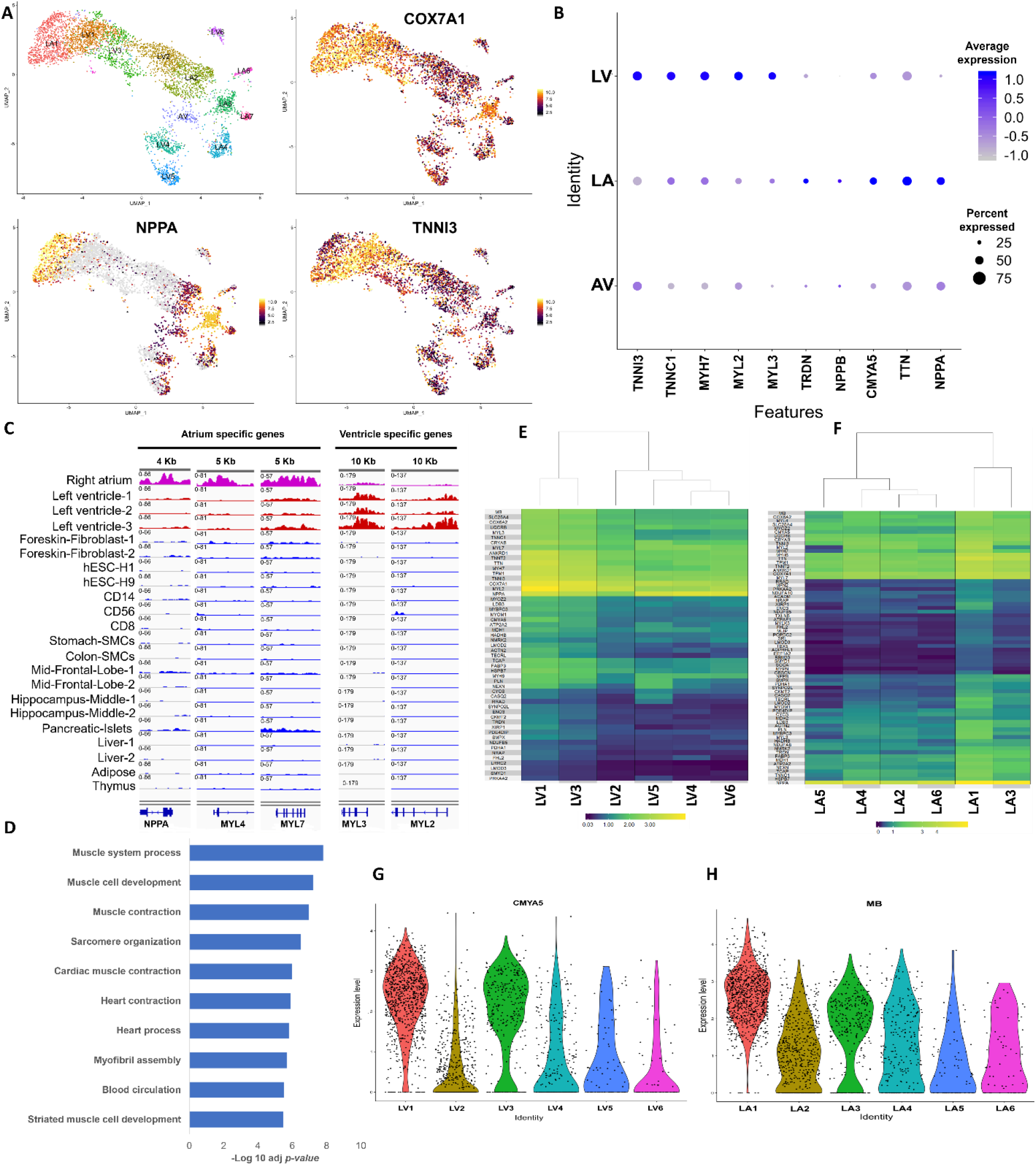
A) UMAP plot indicating the unsupervised clustering of 4,881 CMs from left atrium and left ventricle, which shows there are six left ventricle clusters, six left atrium clusters, and one atrioventricular cluster. COX7A1, NPPA and TNNI1 are represented as genes that have uniform expression levels across all cells, higher expression in the atrium and higher expression in the ventricle respectively. B) Dot plot showing the expression of genes that were differentially expressed between the atrium and the ventricle. For the ventricle, there were four differentially expressed genes compared to the atrium with *p-values* less than 0.049. However, we added TNNC1 (*p-value* = 0.05) to make the graph comparable to the atrium that had 19 differentially expressed genes. The average scaled expression is between −1 and 1, and light blue denotes genes with low expression and dark blue denotes highly expressed genes. The size of the dots indicates the percentage of cells that express the gene. C) H3K27ac peaks on the promotor region of the genes across different cell and tissue types which specifically shows heterogeneity of epigenetic signature between atrium and ventricle. D) Gene ontology of left atrium differentially expressed genes. The terms are ordered based on their −log10 adjusted *p-value*. E & F) Heatmap plots showing heterogeneity of CM-specific genes within the left ventricle (E) and the left atrium (F) clusters. G & H) The violin plots show the cluster label on the x-axis and expression value on the y-axis. Cluster 1 and 3 in both the atrium and the ventricle display the most dispersed gene expression. 53 genes in LV1 differ from the other ventricular clusters, (G) while 73 genes in LA1 differ from the other atrial genes (H). CMYA5 and MB are two examples out of the 118 CM-specific genes.

### Evaluating pluripotent stem cell differentiation toward cardiomyocytes

Differentiating pluripotent stem cells (PSCs) into mature CMs that resemble their *in vivo* counterparts has been a challenge in the field of cardiology research.^23^ Differentiated CMs stray from their natural counterparts in multiple aspects, including transcriptomic features, structure of cytoskeleton and contractile fibers, and metabolic and electrochemical properties.^15^ Because of the rigorous steps we used to obtain a reliable reference of an *in vivo* CM transcriptomic profile in the previous steps, we could compare the success of differentiation protocols from PSCs to this *in vivo* baseline. To this aim, we used the expression data of 15 studies in which CMs had been generated using 10 distinct differentiation protocols (Fig. 4A, Table S3).^24–43^ These protocols use combinations of signaling proteins and small molecules over a range of 7 days to 23 days to make the differentiated CMs. Therefore, the list of compiled protocols covered in our study includes a wide range of differentiation molecules and timelines (Fig. 4A).

**Figure 4.**
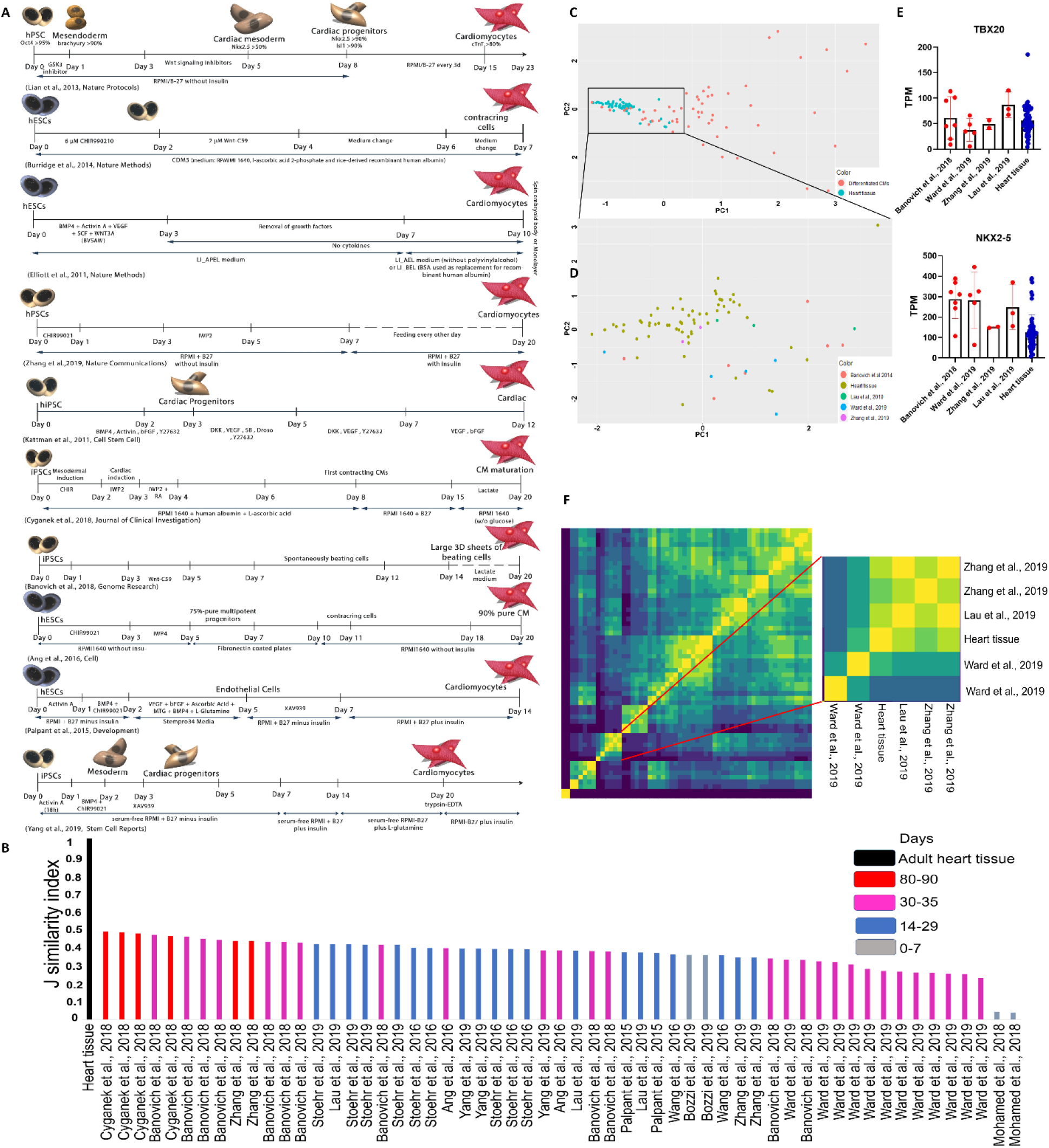
A) An overview of the protocols that are being analyzed in our study. The duration, differentiation molecules added, and the intermediate cells for each protocol are represented. B) Jaccard similarity index which shows resemblance of differentiated CMs to the pooled primary CM samples using expression value all identified genes. The similarity value is represented in the Y axis and different differentiation data setsin the X axis. C) Principle component analysis of the primary heart tissue samples and all differentiated CMs according to the expression of TBX20 and NKX2-5. The heart tissue samples are indicated in green and the differentiated CMs are indicated in red. D) Principle component analysis of the primary heart tissue samples and four studies that result in differentiated CMs most similar to primary heart tissue according to the expression level of TBX20 and NKX2-5. These four studies used two differentiation protocols. Each study is represented with a different color. E)) Bar graphs show the quantified TPM values of TBX20 and NKX2-5 genes in the selected studies and primary heart tissues. Each study is shown as a separate column in red, while the heart samples are represented in blue. F) Jaccard similarity correlation matrix of all bulk pooled human primary CMs as reference to compare with all differentiation data sets. The insert magnifies the pooled primary CMs and the most similar differentiation protocols to the primary CM. Yellow and dark blue colors indicate the highest and lowest similarities between samples.

To evaluate which differentiation protocol generates CMs that are the most similar to those *in vivo*, we calculated the Jaccard similarity index for all genes identified from RNAseq experiments (15349 genes). The range of the Jaccard similarity index is 0 to 1, in which 0 represents the lowest and 1 represents the greatest similarity. Our analysis revealed that studies using Lian et al., 2013 and Burridge et al., 2014 protocols (or their modified versions) produced CMs with the highest similarity to primary CMs (J > 0.4) (Fig. 4B). Furthermore, our results revealed that protocols with longer differentiation periods generally produced CMs more similar to those *in vivo.* For example, ventricular CMs generated from a 90-day differentiation protocol published by Cyganek et al., 2018^31^ (which is a modified protocol of Lian et al., 2013 and Burridge et al., 2014) are the most similar to *in vivo* CMs based on the Jaccard similarity index (Fig. 4B). In addition to considering the entire set of genes which were identified by RNAseq technique, we looked at the expression of the three specific transcription factors previously identified, TBX20, TBX5, and NKX2-5, that play significant roles in the regulation of CM-specific genes during development and differentiation.^44^ Initially, however, we found that TBX5 has a more heterogenous expression in CMs than the other transcription factors and make primary heart samples to be dispersed on Principle Component Analysis (PCA) plot (Fig. S6C). Wang et al., (2020) also reported that TBX5 has a differential expression pattern between the left ventricle and the left atrium.^5^ Therefore, we proceeded with TBX20 and NKX2-5 as common references to evaluate the accuracy of the differentiation protocols. PCA based on expression values of TBX20 and NKX2-5 showed high levels of similarity between primary CMs from all studies, but the differentiated CMs showed high variation between studies (Fig. 4C). Based on the PCA results, we identified four studies that possessed the highest similarity to the primary CMs (Fig. 4D). Three of these studies used two separate differentiation protocols, while the last one used a combination of the two protocols. Studies performed by Lau et al., 2019^43^ and Ward et al., 2019^36^ used a chemically defined differentiation protocol that was introduced by Burridge et al. at 2014^25^. The third study, done by Zhang et al^35^ used a protocol from Lian et al., 2013^24^. Lastly, the Banovich and colleagues study in 2019^28^ used light modified differentiation protocol from both Lian et al., 2013^24^ and Burridge et al., 2014^25^. These four studies exhibit the most similar expression of TBX20 and NKX2-5 to that found in CMs *in vivo (*Fig. 4E). Expressions of these two transcription factors were used to calculate the Jaccard similarity index between pooled primary CMs samples and differentiated CMs, and we found that the Jaccard similarity index (Fig. 4F) showed the same results as identified through analysis of the PCA plots (Fig. 4C, D, and E, Fig. S7A). Notably, Cyganek et al., 2018^31^ and Zhang et al., 2019^35^ protocols were not part of this TF analysis, as these two protocols have longer differentiation time and have higher expression of TBX20 and NKX2-5 (Fig. S7B). Overall, these analyses identified that the two protocols by Lian et al., 2013^24^ and Burridge et al., 2014^25^ produced CMs most similar to those *in viv*o.

### Cardiac muscle cell differentiation protocols fail to generate CM transcriptomes that accurately mirror *in vivo* counterparts

The previous section determined which differentiation protocols produce CMs with the expression of key transcription factors most similar to primary CMs. Here, we further aimed to accurately identify the transcriptomic differences between primary and differentiated CMs. Therefore, we compared expression profiles of all differentiated CMs (57 samples) to that of heart tissue (61 samples). Although two protocols best mimic the production of primary CMs, we did meta-analysis for all differentiation protocols between generated CMs and primary adult CMs to better understand what all of these protocols should focus on in the future. PCA analysis of expressed genes showed that all differentiated samples, regardless of the protocol followed, have distinct gene expression profiles compared to *in vivo* CMs (Fig. S7C). Our meta-analysis revealed 348 genes that were significantly differentially expressed between these groups of CMs (Fig. 5A). 224 of these genes had higher expression in primary CMs, while 124 genes had higher expression levels in differentiated CMs (Table S4). Gene expression clustering showed two distinct clusters, in which all *in vivo* CMs clustered together and all differentiated CMs clustered separately (Fig. 5A). The genes that have higher expression in primary CMs mainly have role in functionality of CMs, like cardiovascular system development and muscle contraction (Fig. 5B and Fig. S8A). On the other hand, 124 genes had higher expression in the differentiated CMs, most of which play a role in cell cycle regulation, cell division, and proliferation (Fig. 5C and Fig. S8B). Portion of genes that have higher expression in primary CMs are among the CM-specific gene list. For instance, MB, COX7A1 and NRAP are genes with very well-known functions in CM that differentiated CMs failed to express at levels comparable to primary CMs (Fig. 5D, Fig S9).^45–47^ The list of genes failed to be expressed by induced CMs is not restricted to those CM-specific genes that we identified in the first step. Instead, there are other well-characterized genes that have crucial roles in CMs development while they also have high expression in other types. For example, FABP4 is involved in the metabolic transition of CMs from glycolysis to fatty acid oxidation, which is an important step in the maturation of CMs.^6^ The failure of induced CMs to express genes like FABP4 adds to the list of difference that they exemplify compared to primary CMs. These genes, as well as others that fail to be expressed by differentiated CMs, are included in Fig. 5E and Fig. S10A. These results indicate that regardless of the differentiation protocol used, differentiated CMs fail to express many genes necessary for cardiac muscle cell function and development. On the other hand, among 124 genes that had higher expression in the differentiated CMs and have role in cell cycle regulation and proliferation we can highlight MYCN gene (Fig. 5F, S8B, Fig. S10B). This transcription factor involved in late embryonic and early postnatal proliferation of CMs and is an example of a gene that must be silenced during normal CM development.^48^

**Figure 5.**
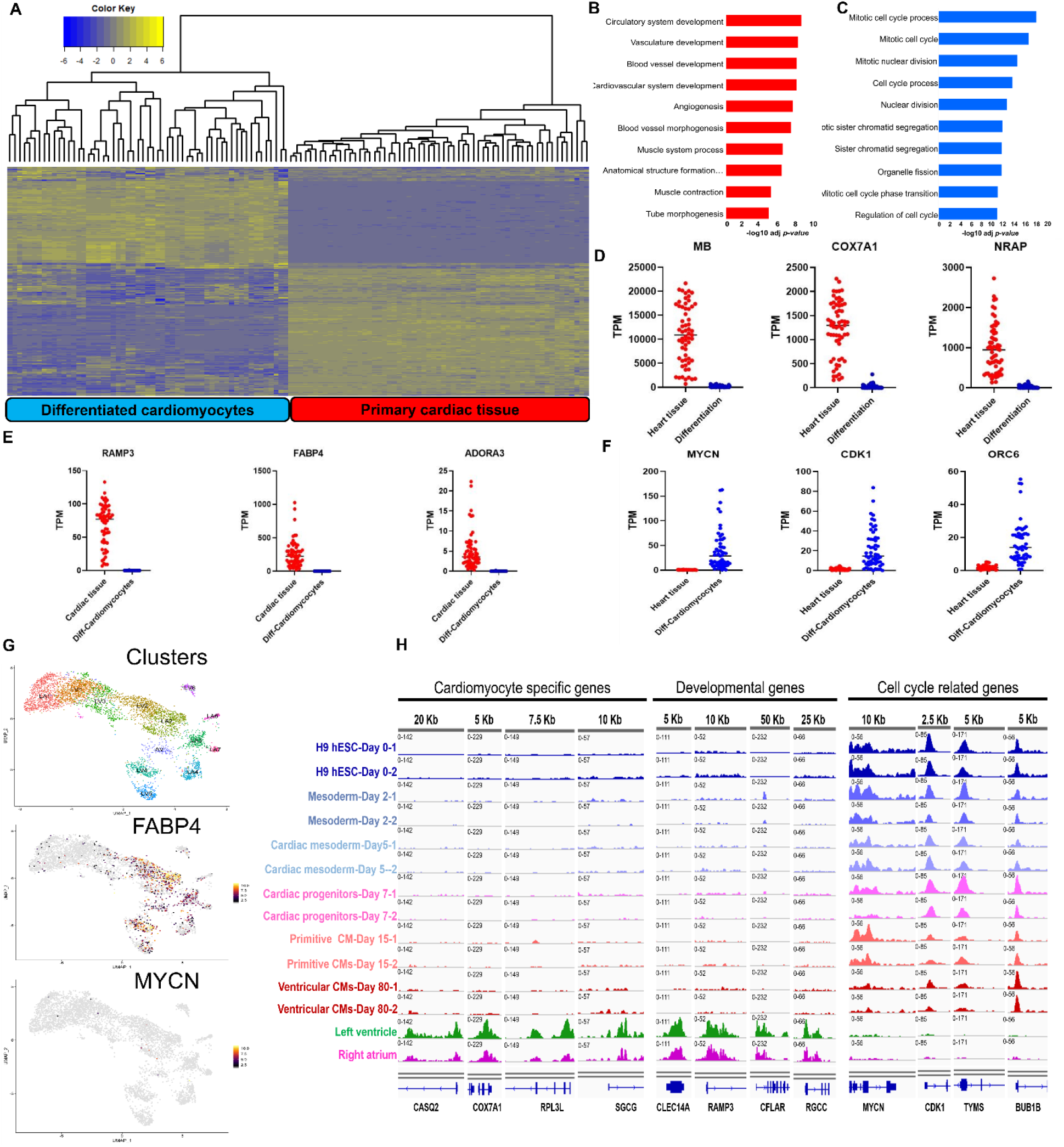
A) Hierarchical clustering of primary heart tissue samples and differentiated CMs according to the expression of genes that were differentially expressed between them. Both primary CMs and differentiated CMs are clustered separately from each other. The up and down regulated genes are represented in yellow and blue respectively. B & C) Gene ontology analysis of the genes that have higher (B) or lower (C) expression in primary CMs compared to differentiated CMs. The terms are ordered according to the −log10 *p-value*. D-F) Expression plots that quantify the TPM values of genes with differential expression in primary versus differentiated CMs. (D) graphs the genes with absolute highest expression in the heart tissue and are among the list of 118 CM-specific genes. (E) plots genes with higher expression in primary CMs not restricted to the list of 118 genes, while (F) plots genes with higher expression in differentiated CMs. G) Upper panel: UMAP plot displaying the distribution of atrium and ventricle clusters in the heart tissue. Middle panel and lower panel: UMAP expression plot of FABP4 and MYCN. The yellow and grey colors represent higher and lower expression of genes respectively. H) we assessed H3K27ac statutes in promoter region of CM-specific genes, genes involved in cardiovascular system development and cell cycle related genes. Different differentiation time points were compared with adult H3K27ac ChIP-seq derived from ventricle and atrium..

The above data was generated from bulk RNA-seq datasets. Because some of the gene expression data may therefore come from non-CM cells like smooth muscle cells, fibroblasts, and endothelial cells, we cross-referenced our data to the single cell RNA-seq data from Wang et al (2020).^5^ This data confirmed that many genes that are not among the CM-specific profile, like FABP4 and genes involved in cardiovascular system development, are expressed in primary CMs (Fig. 5G and Fig. S10A). In addition, single cell gene expression analysis showed that 124 genes present in differentiated CMs have lower expression or are not detectable in primary CMs, including MYCN, CDK1, CDC20, ORC6, TOP2A, BUB1B and AURKB (Fig. 5F and S10B). In summary, we showed that differentiated CMs fail to establish a gene expression profile that fully resembles *in vivo* human adult CMs and instead retain a more progenitor-like transcriptome.

Finally, to check the chromatin accessibility of the differentiated CMs and compare them with the chromatin state of the primary CMs, we used the ChIP-seq data for H3K27ac generated by Zhang et al., 2019 and human epigenome roadmap.^22,35^ Zhang et al., 2019 data has two features that make it a reliable source of information for further analysis. First, this paper uses a Lian et al., 2013 differentiation protocol, which we previously identified as producing the CMs most similar to *in vivo* CMs. Secondly, they retain the cells in culture for longer to generate ventricular CMs (about 80 days), which again we demonstrated as increasing the similarity between induced and primary CMs. Therefore, the epigenetic data produced by this protocol possesses both features and makes it ideal for further analysis. We looked into the status of H3K27ac marks during different stages of CM differentiation compared to adult human heart tissue. We first investigated the acetylation pattern of CM-specific genes that were not expressed in the transcriptome data of the differentiated CMs. As expected, we found that genes which were not expressed by differentiated CMs similarly lacked H3K27ac marks (Fig. 5H). These genes included CASQ2, COX7A1, RPL3L, and SGCG, all of which lacked H3K27ac and mRNA expression in differentiated cells but were present in primary CMs. Regardless of the duration of the differentiation protocol, none of the differentiation durations resulted in H3K27ac marks for these genes (Fig. 5H). While these CM-specific genes failed to be acetylated in differentiated CMs at all time points, acetylation status of a group of genes were indeed dependent on the differentiation time. Differentiation of ventricular CMs for 80 days resulted in the similar acetylation pattern of CM-specific genes like COX6A2, NRAP, LMOD3 or genes involved in cardiovascular development like CAV2 between induced and primary cells (Fig. S11). These genes acquired the same H3K27ac pattern at day 80 of differentiation but not at the earlier stages of differentiation (Fig. S11). Other genes that have important roles in CM development, but which are not restricted to the list of CM-specific genes, also have different acetylation pattern in induced CMs compared to primary CMs (Fig. 5H). For example, all differentiation protocols failed to result in RAMP3 H3K27ac marks, while CMs from the left ventricle and right atrium have H3K27ac marks in their promoters.

We also investigated the levels of cell cycle-related genes, including MYCN, CDK1, BUB1B, and TYMS and found that differentiated CMs have higher expression of these genes. Consistent with their increased expression, differentiated CM had increased H3K27ac marks, similar to pluripotent stem cells, on these cell cycle-related genes (Fig. 5H). Notably, the *in vivo* CMs lack both the mRNA expression and acetylation of these genes. Therefore, there is a significant correlation between continued expression of these genes and their active epigenetic marks. An inability to silence these genes in differentiated CMs may provide one explanation for the discrepancy observed between primary and differentiated CMs. In other words, *in vivo* CMs regulate their cell cycle status, while differentiated CMs may be unable to exit the cell cycle and thus exhibit more progenitor characteristics.

In this study, we analyzed an extensive set of transcriptomic data to define the specific transcriptomic landscape of primary and differentiated CMs. We identified 118 genes that are highly expressed in CMs compared to all other cell types (Fig. 6). These genes are critical for CM function and development, including cardiac muscle fiber organization, energy metabolism, and cardiac development and function (Fig. 6). 19 of these genes showed spatial gene expression heterogeneity between the left ventricle and the left atrium. By using the expression profile of the master transcription factors of CM development, we then assessed available PSC to CM differentiation protocols to find the protocols that generate CMs most similar to adult human CMs, with two protocols determined to be the most successful. Finally, upon comparing the full transcriptome of differentiated CMs to primary CMs, we found that all differentiation protocols result in CMs that retain more embryonic and proliferative features than do primary adult CMs. Overall, our study provides novel insight into the specific transcriptomic landscape of CMs, including confirming the spatial heterogeneity between CMs in different parts of the heart. Furthermore, we demonstrated the specific weaknesses that all differentiation protocols exhibit, which is a critical step in determining the future changes needed to generate CMs that are suitable for regenerative medicine and research.

**Figure 6.**
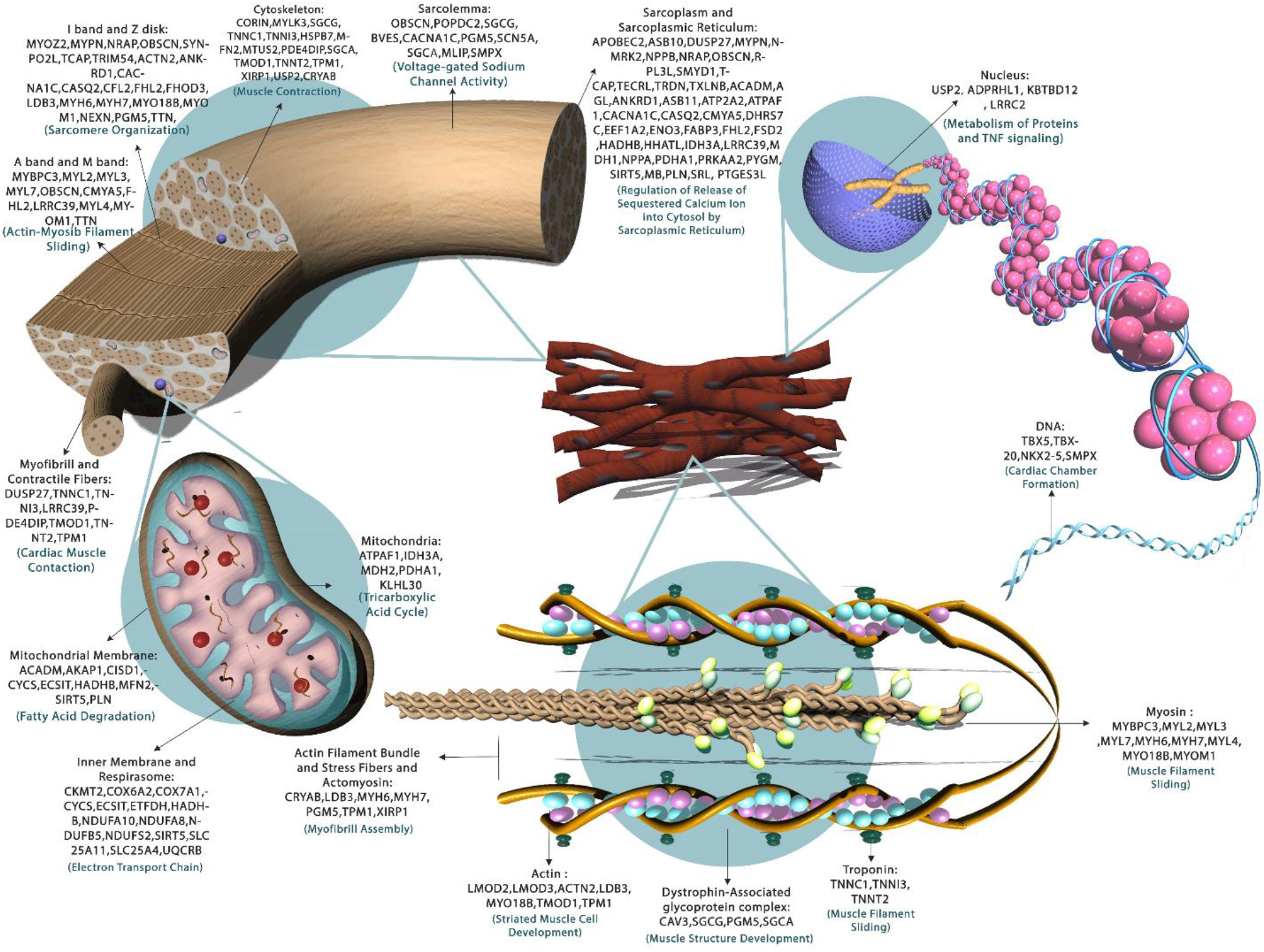
Schematic diagram of the cellular location and molecular function of the identified 118 CM-specific genes. The identified genes have various functions, including controlling expression of the genes, as transcription factors, cytoskeleton formation, mitochondrial function, and formation of skeletal bundles.

## Discussion

Transcriptomic analysis is a powerful tool used to understand the identity and function of cells. In the present study, we focused on unraveling the unique transcriptomic profile of human and mouse primary CMs that subsequently allowed for the comparison of primary CMs to differentiated CMs. We first gathered the transcriptomic information of 91 cell types and subtypes. Danielsson and colleagues showed that integrating RPKM/FPKM values from datasets generated in different laboratories with different protocols successfully results in the clustering of tissues based on the tissue type and not on the laboratory of origin.^49^ The same group found that using FASTQ file to generate FPKM values had no major benefit compared to using the RPKM/FPKM values.^49^ Additionally, the use of different analysis pipelines on RNA-seq datasets has been shown to yield 88% similar results for protein coding genes.^50^ In our study, we converted the FPKM/RPKM values from multiple datasets to TPM values, which homogenously mixed the data for further analysis. We then used a single analytical technique to determine the differentially expressed genes. Although we did not use FASTQ files, our large sample number provided reliable and valid statistical results. In addition, we found a strong correlation of differentially expressed genes between our dataset results and the human protein atlas (HPA) datasets. Furthermore, single cell RNA-seq data and different ChIP-seq data sets confirmed our transcriptomic results. Following comprehensive data mining, provided a list of genes that have higher expression in mouse and human CMs compared to all other cell types. While our study investigated a large and diverse number of cell types, one limitation of the current study is that we could not include transcriptomic analysis of every somatic cell type that exists in the body. This would have increased the power of our findings and may have heightened the specificity of genes found to be expressed by CMs. For example, we have 61 heart tissue samples and 34 smooth muscle cell samples, but we have no samples specific to skeletal muscle. Using datasets from the (HPA), we discovered that a portion of the 118 genes identified as CM-specific genes indeed have high expression in skeletal muscle tissues.^21^ Even this comparison between our findings and data from the HPA dataset is imperfect, as their data contains signals from various cell types present in a single tissue. Because we compared our data to well-defined, single cell CM data, we are confident that our results are accurate for detailing the expression profile of CMs compared to most other cell types. To increase the validity of H3K27ac of our findings, we used human epigenomic data from ChIP sequencing experiments of H3K27ac mark. Indeed, our transcriptional findings of primary and induced CMs were in agreement with the epigenetic signature of the genes. The project would be more impactful if it included proteomic data, which could have been further supplemented with functional confirmation. Despite this possible addition that could increase the validity of our findings, transcriptomic expression profiling coupled with epigenetic confirmation has been proven to be a reliable source of information that can lead to greater conclusions and benefit both research and medicine.^21,22,51^ Additionally, our results provide important guidance for future proteomic and functional studies, as we have identified specific genes and pathways implicated in these cells.

As expected, the majority of the genes that were upregulated specifically in CMs were genes with well-known roles in CM function and development. For example, master regulators of CMs, including NKX2-5 and TBX20, were among the identified upregulated genes.^52,53^ In addition to the expected genes, we also found a small group of genes specifically upregulated that do not have an identified role in CMs, such as HHATL. This gene might have a role in the formation of a septum between ventricles as it is involved in the sonic hedgehog (SHH) signaling cascade.^54^ Obtaining a list of CM-specific genes, as we have done here, may have a number of applications. For instance, it could potentially be used in regenerative medicine to check the extent of CM differentiation based on the similarity to their *in vivo* counterparts. This list may also act as a reliable source for characterizing primary CMs during development.

The majority of studies that have profiled the transcriptome of CMs have performed their investigations on the entire heart tissue, even though the heart at least contains five different cell types.^5^ In our study, although we obtained the list of CM-specific genes using microarray and bulk RNA-seq techniques, we confirmed our results using publicly available heart single cell RNA-seq datasets. By doing this, we were able to benefit from the advantages of each technique. We had the large cell number and experiment sizes from bulk RNA-seq, and we could compare this data to the very specific, though small power samples from single cell RNA-seq. Through this approach, we identified CM-specific genes and were able to identify spatial heterogeneity present in the heart tissue. For instance, we confirmed that NPPA has increased expression in mouse and human primary CMs compared to other somatic cells, and we noted that its expression is specifically increased in the atrium compared to the ventricle. It has previously been indicated that NPPA is reduced in the ventricles upon birth.^55^ Furthermore, it has been shown that NPPA mutations are correlated with diseases, such as atrial fibrosis.^56^ Moreover, it has been indicated that NKX2-5 and TBX20 bind to the NPPA promotor and control its expression.^57,58^ In our study, we confirmed the expression of these genes specifically in CMs and used ChIP sequencing data to confirm a direct interaction between TBX20 and NPPA. Altogether, with the expression of key transcription factors and known CM-specific genes that agrees with previous literature, this is the most thorough and high-throughput study of the CM transcriptomic profile to date.

Another aim of this study was to clarify the differences between primary and differentiated CMs based on their transcriptional profiles. To this aim, we compared the gene expression profiles of primary and induced CMs and concluded that the populations differed in two ways. Firstly, the induced CMs fail to express genes critical for development and function of CMs. For example, induced CMs do not express COX7A1, the absence of which results in dilated cardiomyopathy in mouse studies.^46^ Furthermore, NRAP is the second largest actin-binding cytoskeletal protein and is expressed in myofibril precursor cells during myofibrillogenesis.^59^ Homozygous truncated NRAP genes are found in the whole exome sequencing data of dilated cardiomyopathy patients which highlights the importance of this gene CMs development.^47^ In our analysis, we confirmed that induced CMs have lower expression of this gene compared to primary CMs (Fig. 5E). Secondly, the induced CMs aberrantly down regulate positive regulators of the cell cycle, preventing them from becoming fully differentiated (Fig. S5B and Fig. S7B). For instance, MYCN, which is known to induce cell cycle progression and proliferation, is more highly expressed in differentiated cells than in primary adult CMs.^48^ It has been shown that MYCN is regulated by Sema6D that regulates cardiac muscle cell proliferation and prevents differentiation of cells.^48^ Therefore, differentiation protocols may aim to down-regulate MYCN to inhibit proliferation and promote differentiation of CMs that would make them more similar to their primary counterparts. Altogether, these findings provide a detailed view of the current state of CM differentiation protocols compared to *in vivo* CMs, and it may guide future research that will produce more impactful CMs.

To the best of our knowledge, this is the first study that has compiled such an extensive and comprehensive transcriptomic profile of primary CMs. The large number of cell types and subtypes investigated, combined with the high consistency between our findings and those in the HPA dataset, make our findings very reliable. Based on this data, the detailed primary CM transcriptomic profile may guide future CM differentiation protocols that were analyzed in this same study.

## Methods

### Data collection and preprocessing

#### RNA-seq data

Through careful mining of the Gene Expression Omnibus (GEO), we gathered a comprehensive and unique repertoire of mouse and human RNA sequencing data for healthy (normal/wild type) somatic and stem cells. The downloaded RNA-seq datasets contained read counts, reads/fragments per kilobase million (RPKM/FPKM) and transcript per million (TPM) values. Samples with low sequencing depth or few detectable genes (less than 22000 genes for somatic cells and stem cells) were excluded as low-quality samples before any further analysis. After excluding these samples, we had 87 data sets with 626 RNA-seq samples from either humans or mice. Read counts were normalized based on the gene length and number of aligned reads to the reference genome, generating TPM values. To obtain TPM from read counts, first we converted read count values to FPKM/RPKM for each gene. To this end we used following steps:

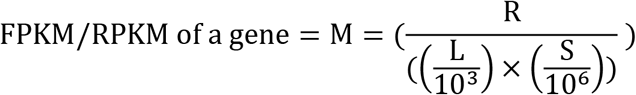

Where each symbol is equal with following factors

R: Total number of reads mapped to a given gene

L: Length in base pair of a given gene

S: Total aligned read In the next step, following equation was used to calculate TPM from calculated RPKM/FPKM:

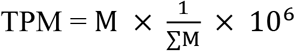

where

M = FPKM/RPKM value for a gene

Σ M = The sum of FPKM/RPKM values of a given library. ^60,61^

To detect outliers in the generated source of normal human and mice cells we used TPM values for PCA analysis and hierarchical clustering. TPM values were used to perform PCA using the built-in function of R called *prcomp*, and the results were visualized for 2D and 3D PCA using the ggplot2 and PCA3D packages in R, respectively. The first and second principle components were used to remove outliers. Overall, 57 RNA-seq samples were removed from 15 independent datasets (studies), resulting in 584 RNA-seq samples that we continued to use (Table S5).

In addition to bulk RNA-sequencing, single cell RNA-seq datasets GSE109816 and GSE121893 from Wang et al., (2020) were used to investigate CMs more specifically. Single cell RNA-seq was also used to explore the gene expression heterogeneity of cells.^5^

Microarray data sets from 62 studies with 273 samples were also used to analyze the transcriptome of cells. We downloaded affymetrix mouse genome 430 2.0 array raw files (.CEL files). All samples were filtered to only include normal somatic cells (Table S5). In addition, all data was normalized using Robust Multichip Average (RMA) in R through the AffylmGUI package.^62,63^

Finally, to add further confirmation to the transcriptomic analysis, we analyzed H3K27ac ChIP-seq data from the human epigenome roadmap and Zhang et al., (2019) datasets ^22,35^ From the human epigenome roadmap dataset, we acquired ChIP-seq data for the right and left ventricle and the right atrium. Additionally, we downloaded data from 14 different cell and tissue types, including foreskin fibroblasts, H1 embryonic stem cells, H9 embryonic stem cells, CD8 cells, CD14 cells, CD56 cells, stomach smooth muscle, colon smooth muscle, mid frontal lobe, hippocampus, pancreatic islet cells, liver, adipocytes, and thymus. All samples were download in wig format and were converted into TDF format using igvtools. They were visualized using Integrative Genomic Viewer (IGV)^64,65^ We also used H3K27ac data from Zhang’s et al., 2019, which contains different differentiation time points of H9 human pluripotent stem cells as they were driven towards CMs. These included day 0 (embryonic stem cells), day 2 (mesoderm), day 5 (cardiac mesoderm), day 7 (cardiac progenitor cells), day 15 (primitive CMs), and day 80 (ventricular CMs).^35^

### Meta-analysis

To get a specific transcriptomic profile of CMs, we compared the gene expression profile of CMs with all cells compiled in our source. To do this with a meta-analysis approach, we created a transcriptomic reference of CMs and subsequently compared all other somatic and stem cell types to this reference. These transcriptomic references were made for both human and mouse CMs’ RNA-seq and microarray samples separately (Fig. S12A). In each category of mice and human RNA-seq and microarray data we obtained DEGs (Fig. 12). For example, to make this reference for human RNA-seq expression datasets, we merged 61 human heart tissue RNA-seq samples from three independent studies and considered them as a single reference to compare against all human somatic and stem cell RNA-seq samples. In order to handle such enormous amounts of data, NetworkAnalyst^66^ was used was used, but it has a limitation that prevents analysis of all datasets at the same time. To overcome this obstacle, we compared the expression profile of CMs to five and six somatic/stem cell datasets at a time for human and mouse respectively, which resulted in 16 groups and 81 comparisons for human RNA-seq samples and 3 groups and 18 comparisons for mouse RNA-seq samples (Fig. S12B).

To prepare the data for meta-analysis, we inputted our values into the “multiple gene expression table” section of NetworkAnalyst. After uploading the data as a text file, we converted the gene IDs to “Official Gene Symbol”, thus creating a common ID format for all of our data. We used the “visualization data plot” to check the distribution of values within samples, and we performed differential expression analysis on the data using *p-value* of 0.05 as a cut-off, which was generated using Benjamini-Hochberg’s False Discovery Rate. For meta-analysis, the combined effect size, combined *p-value*, vote counting, and combined fold change (direct merging) were the four criteria used to detect significantly differentially expressed genes (DEGs), though the combined effect size and combined *p-value* were the primary criteria used to select the DEGs (Fig. S12). The effect size considers the size of the difference between two groups and is calculated by dividing the mean differences of the comparisons by the standard deviation. Based on the Cochran’s Q tests we used random effects model to determine effect sizes (genes with *p-value* < 0.05 were kept as significant genes). To calculate combined *p-values* we used Fisher’s method, which is a weight-free method (combined *p-values* < 0.05 were kept as significant genes). Vote counting is a way to identify genes that are significant across all comparisons during meta-analysis. For vote counting we kept genes with vote counts 5 and for combined fold changes, we kept genes with *p-values* <0.05 and fold change >2 as significant genes. For each criterion, we considered only significant genes that passed a certain threshold (Table S6). We then extracted genes that were significantly differentially expressed as measured by all four criteria, thus removing any genes that were deemed non-significant by one of the metrics. Finally, we merged all genes from the human and mouse RNA-seq and microarray datasets to create a finalized list of significantly differentially expressed genes in CMs compared to all other somatic and stem cells in our datasets (Fig. S12).

### Single cell RNA-seq data analysis

All single cell analysis was performed with version 3 of Seurat.^67,68^ The raw umi matrix of the heart single cell study was downloaded from GEO with GSE109816 and GSE121893 accession numbers.^5^ We excluded the cells that expressed more than 72 percent of mitochondrial genes, which was suggested and performed in Wang et al, (2020). Following data normalization using a global-scaling normalization method called “LogNormalize”, the top 2000 highly variable genes were identified and scaled. This scaled matrix was then the input file for principle component (PC) analysis. Elbow plot were used to determine the number of PCs for down stream analysis. These PCs were used to compute the distance metric, which then generated cell clusters. Unsupervised clustering was done with original Louvain algorithm using *FindNeighbors* and *FindCluster* functions in Seurat. Resolution 0.3 to 0.8 were tested and according to the expression of marker genes the best resolution than clearly distinguish different clusters were selected for further analysis. Uniform manifold approximation and projection (UMAP) was used to visualize clustering results. Differentially expressed genes (DEGs) were found using the *FindAllMarkers* (or *FindMarkers)* function that ran Wilcoxon rank sum tests. Genes with *p-value* less than 0.05 and log2 fold change > 0.25 were considered as DEGs consistent with Seurat package recommendation.^67,68^

### Network analysis

To construct a gene regulatory network, transcription factor (TF)-binding sites were obtained from the Chromatin Immuno Precipitation (ChIP) Enrichment analysis (ChEA) database (Chen et al., 2013; Kuleshov et al., 2016).^69,70^ ChEA contains data from ChIP experiments that determine TF-DNA interactions.^69,70^ We input DEGs and retrieved the TFs that significantly (*p-value* <0.05) regulate those DEGs above a fold change of 2. There are three significant ChIP-seq experiments for TBX20 (from heart mouse), NKX2-5 (HL-1 Cardiac Muscle Cell Line) and TBX5 (HL-1 Cardiac Muscle Cell Line). Using differentially expressed TFs (DE-TFs) and DEGs, gene regulatory networks (GRNs) were constructed using Cytoscape.^71^ CentisCape, a plugin accessed through Cytoscape was used to identify the central TFs that regulated the majority of DEGs.^72^ CentisCape considers the number of direct interactions between a TF and its targets, with this value quantified as degrees. The greater the number of degrees, the more central a TF is.

Protein-protein interactions (PPIs) for DE-TFs were obtained from the STRING database for analysis in our dataset.^73^ These interactions were input into Cytoscape to create a PPI network, as explained in other manuscripts.^74^ This network was analyzed by the Cytoscape MCODE plugin.^75^ For our analysis, complexes that have score greater than 2.0 were considered significant protein complexes.

g:Profiler was used for gene ontology analysis, and results were filtered based on the adjusted-*p-value*.^76^ TPM values from RNA-seq experiments were used to generate heatmaps using ggplot2 and Superheat packagesuperheat packages in R.^77^

### Jaccard similarity matrix calculation

The weighted Jaccard similarity index calculates the similarity between two numeric vectors (in our study two RNAseq samples). The range of this similarity is from 0 to 1, in which 1 is highest similarity and 0 is lowest similarity between samples. The Jaccard similarity index is calculated using the sum of minimum values in both vectors (the intersection of two RNA-seq data samples) divided by the sum of the maximum values in both vectors (the union of each). We used MATLAB code to calculate the Jaccard similarity index automatically.

## Author contributions

Design the experiment and analyze the data: MO, ES, NK, SJE, HH, MY and AM. Writing and editing the paper NK, NWK, MY, AM. Making the figure NK, NWK, SJE, MO, AM.

## Supplementary legend

### Supplementary tables

Table S1. Expression of canonical CM markers genes in the gathered heart tissue datasets. The TPM values are represented in the table.

Table S2. The TPM value of 118 genes across all human RNA-seq samples which were identified in this study as CM-specific expression profile.

Table S3. The list of differentiation data set and differentiation protocols that we used in this study.

Table S4. The list of 348 genes that differentially expressed between primary and induced CM including 224 up and 124 down regulated genes in primary vs induced CMs.

Table S5. Detailed information of all bulk RNA-seq and microarray samples have been used in this study.

Table S6. Combined effect size, combined *p-value*, vote counting and direct merging were used to identify differentially expressed genes.

### Supplementary figures

**Fig S1.**
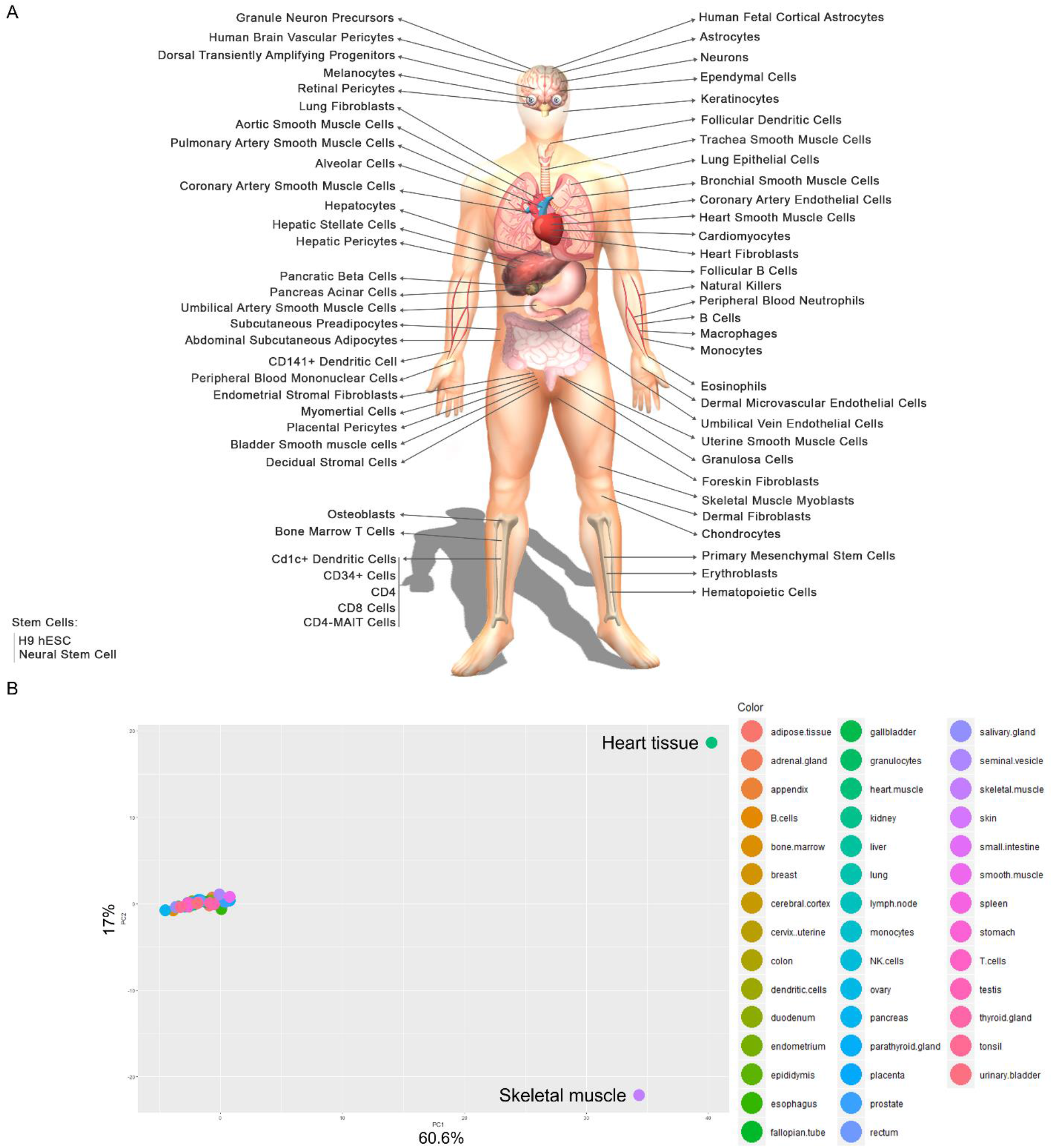
A) Schematic representation of the human cells and tissue that is used in this study. B) Principle Component Analysis (PCA) using expression of 118 CM specific genes which clearly separate CMs from skeletal muscle and other somatic cells.

**Fig S2.**
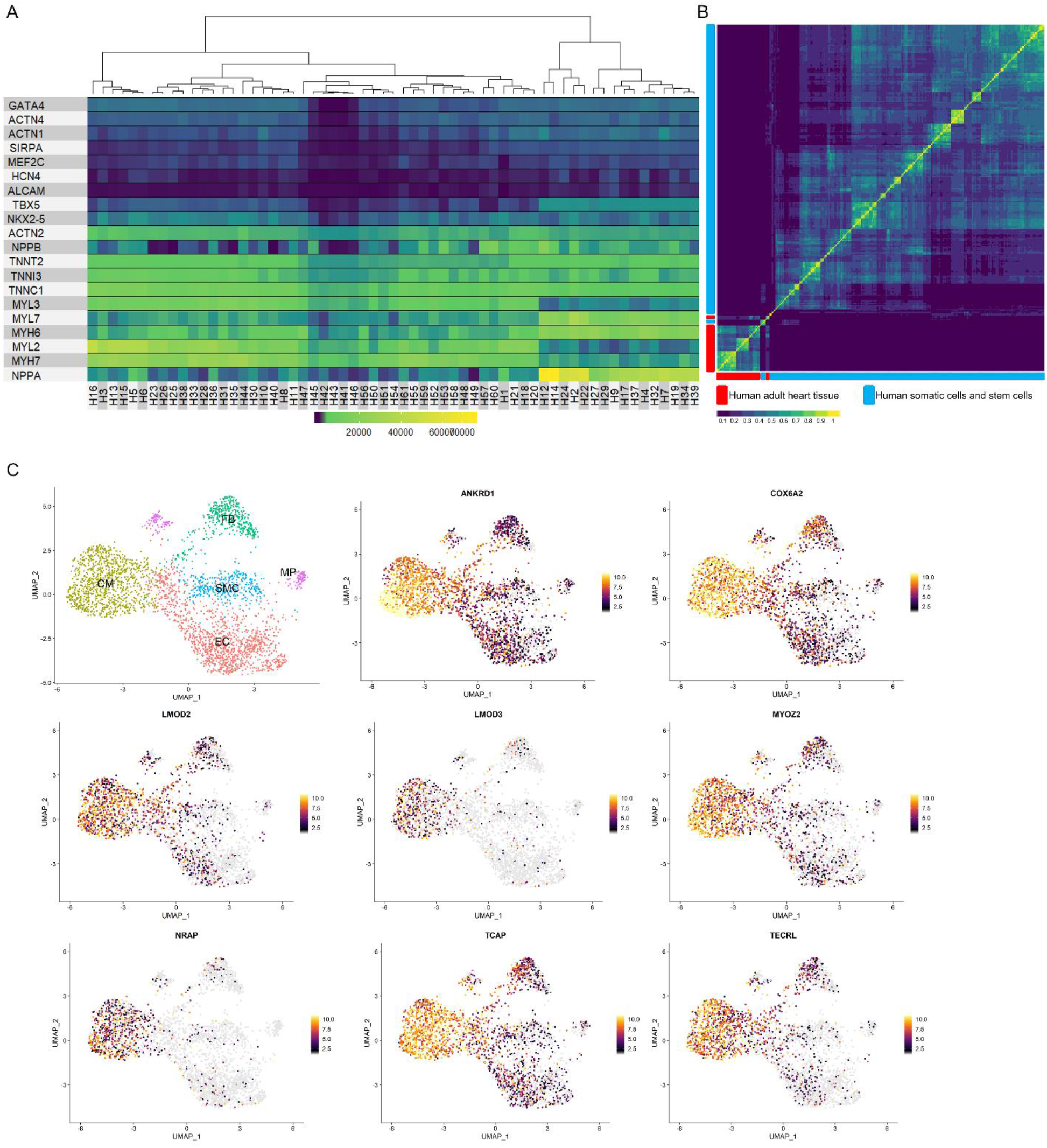
A) The heatmap representing expression of CMs canonical genes expression in the 61 heart tissues. B) Jaccard similarity correlation matrix of all bulk RNA seq datasets. Human adult heart tissues are clustered together and separate from other cell types. Yellow and dark blue colors correspond to similarity index 1 and 0 respectively, indicating the highest and lowest similarities between samples. C) Expression of the eight CM-specific genes are represented on the UMAP plot of the heart tissue. The expression of genes is indicated by a color gradient in which yellow show high and grey low expression.

**Fig S3.**
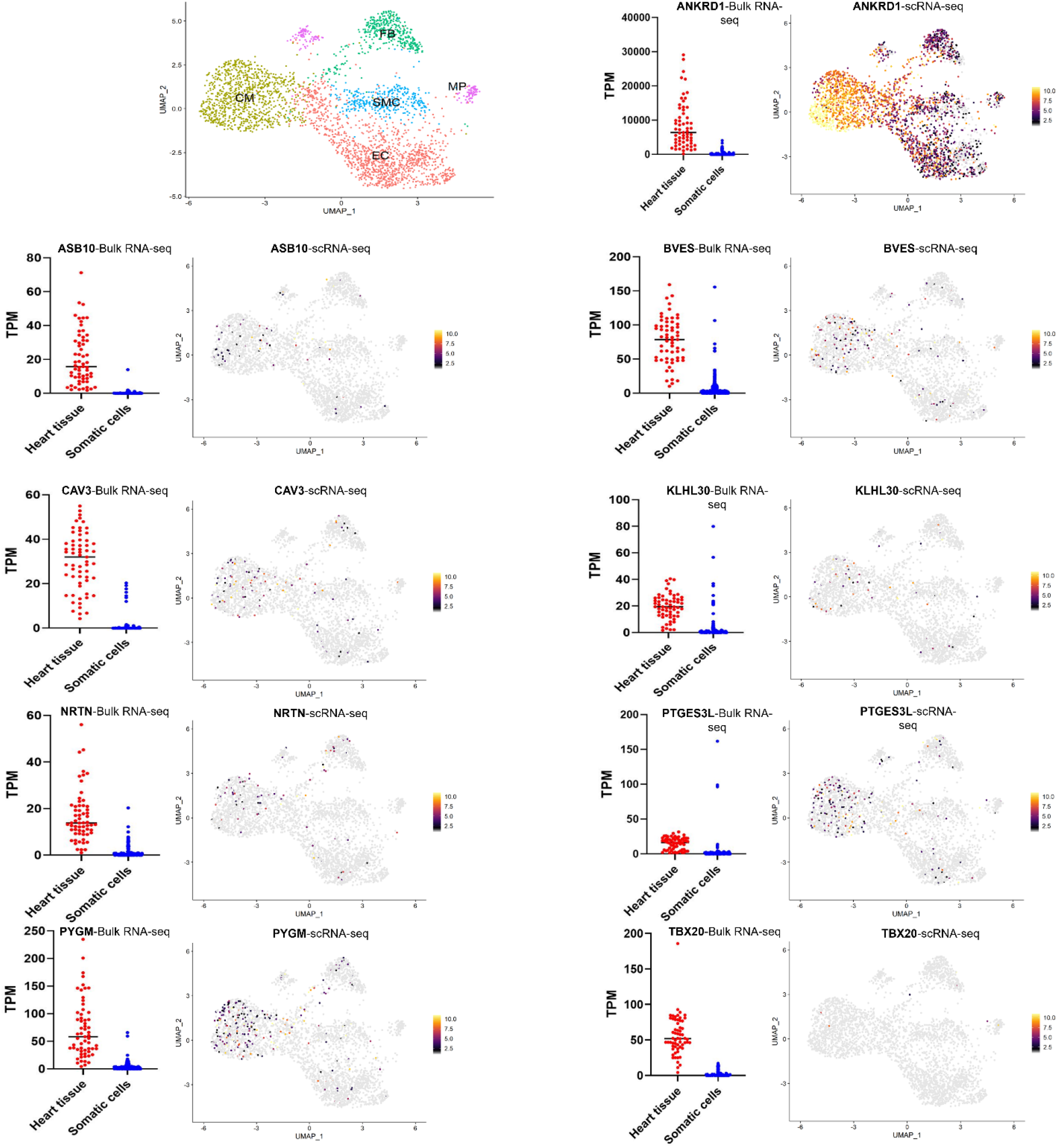
Expression of the lowly expressed genes is represented by bulk RNA-seq and the TPM values are used to plot the graph. Due to the low sensitivity of single cell RNA-seq, this technique could not detect their expression. ANKRD1 is represented as an example of highly expressed genes for comparison.

**Fig S4.**
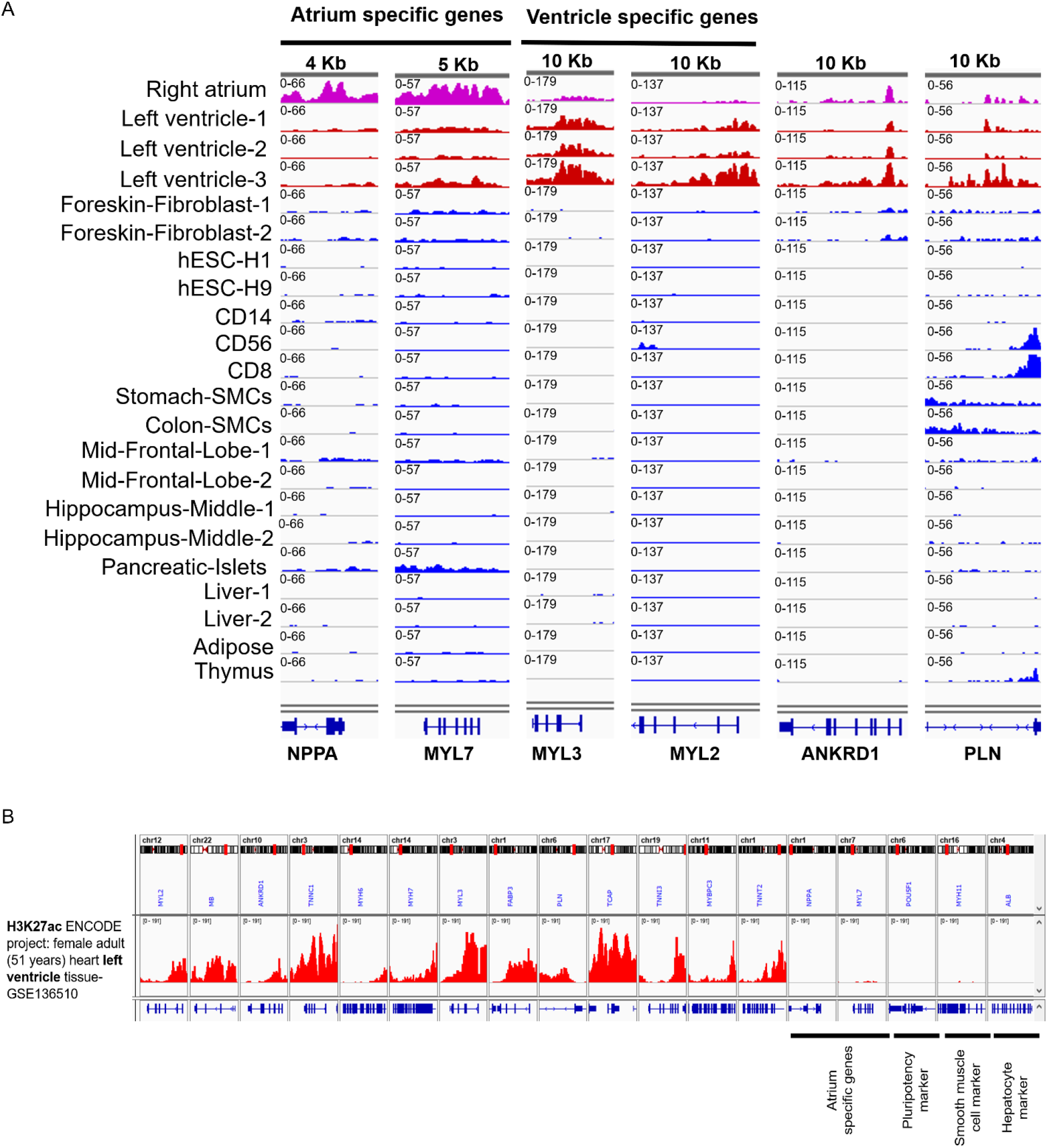
A) Representation of H3K27ac marks for some of atrium and ventricle specific genes across heart and 14 other cell types. B) H3K27ac ChIP-seq experiment from left ventricle of adult female downloaded from ENCODE project. NPPA and MYL7 are atrium specific genes and the samples that is being analyzed in this panel is ventricle sample, so these two genes does not have H3K27ac peak, however, in the Fig. S4A their H3K27ac pattern is obvious in the right atrium row.

**Fig S5.**
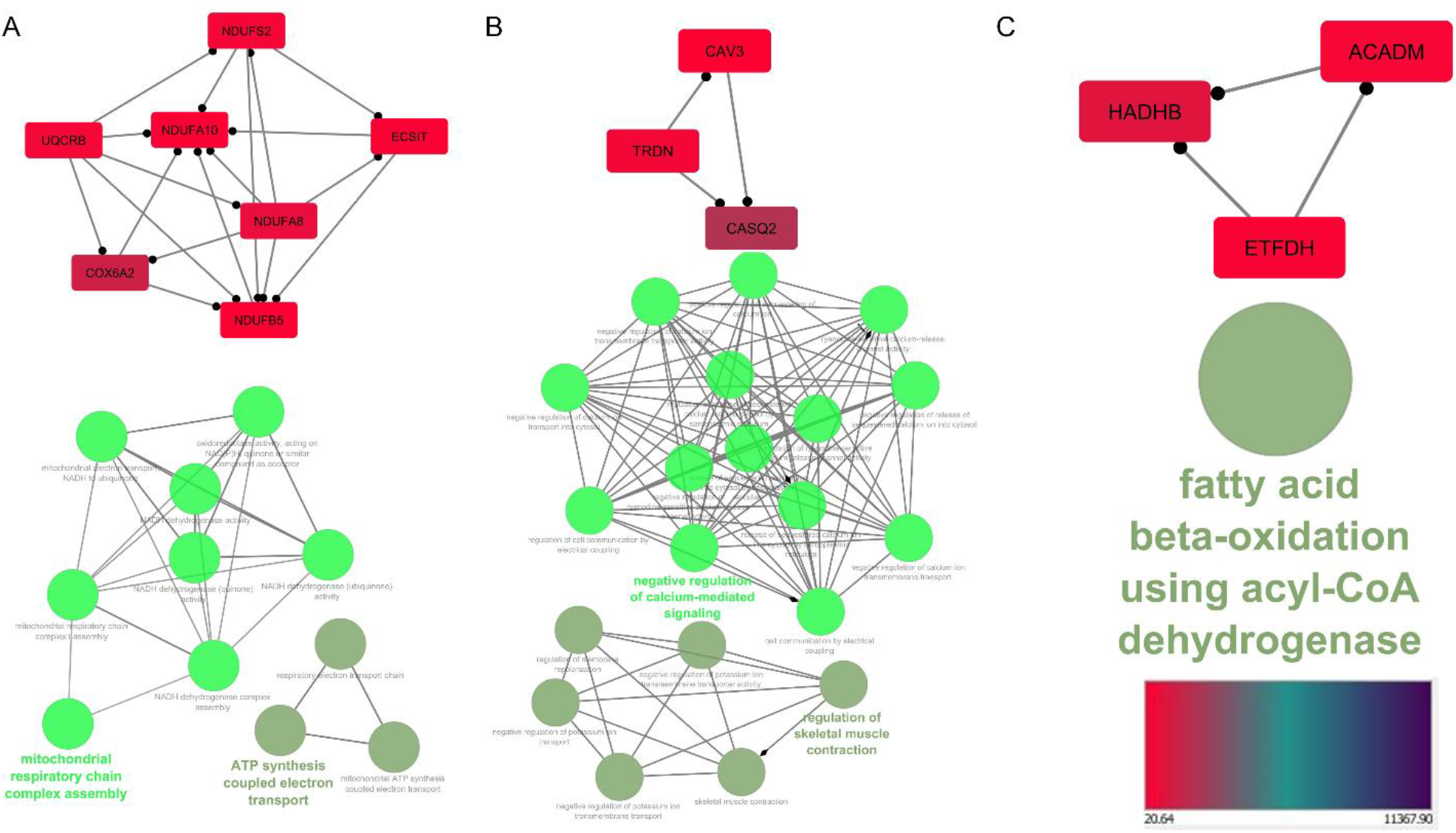
A-C) significant protein complexes of the gene regulatory network and their ontology analysis. Each complex and its ontology is represented in a separate panel.

**Fig S6.**
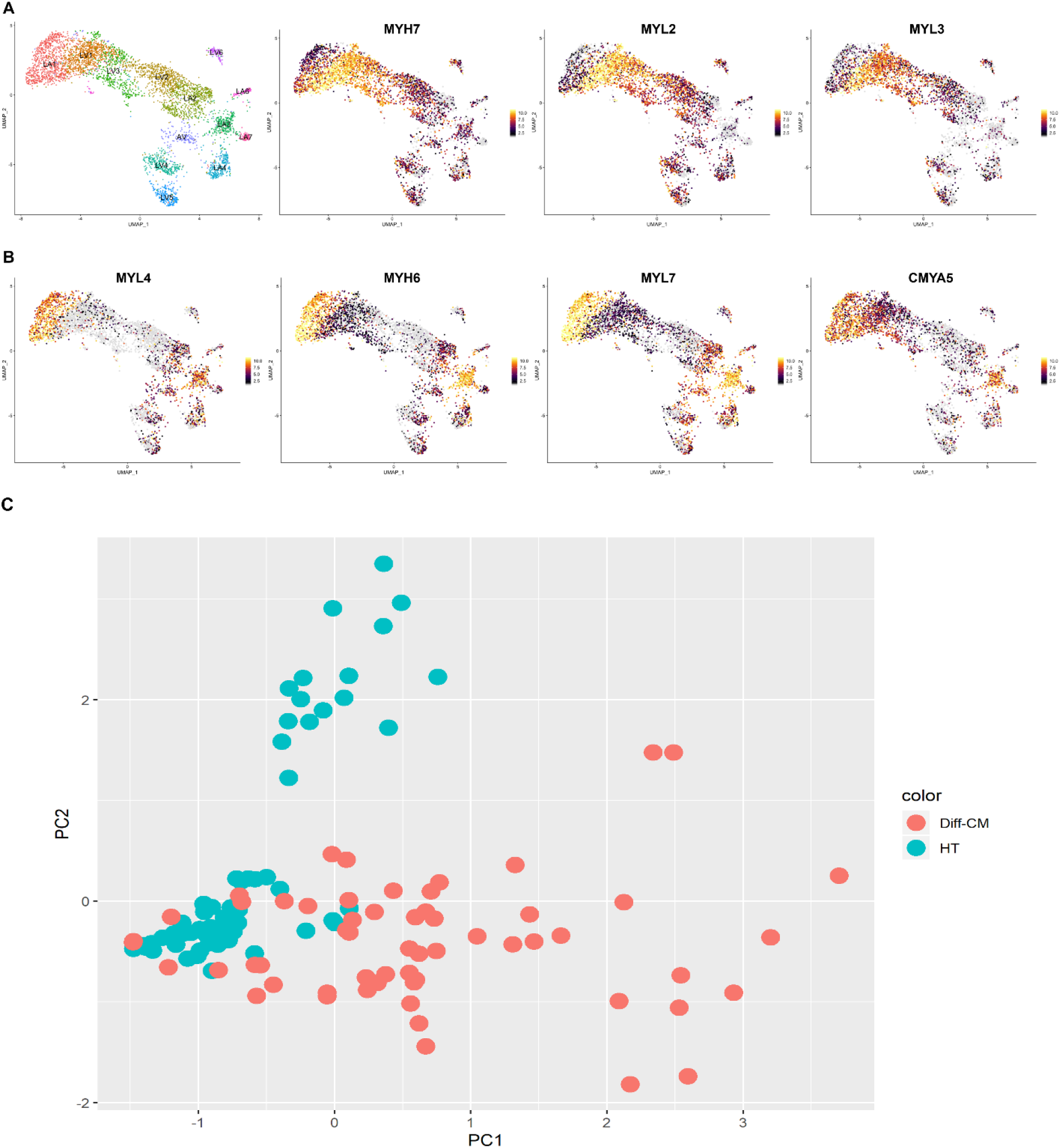
A) The expression of three out of four ventricle specific genes B) The expression of four out of 15 atrium specific genes. C) The principle component analysis of the primary heart CMs and differentiated CMs according to the expression of the three transcription factors including TBX5, TBX20 and NKX2-5.

**Fig S7.**
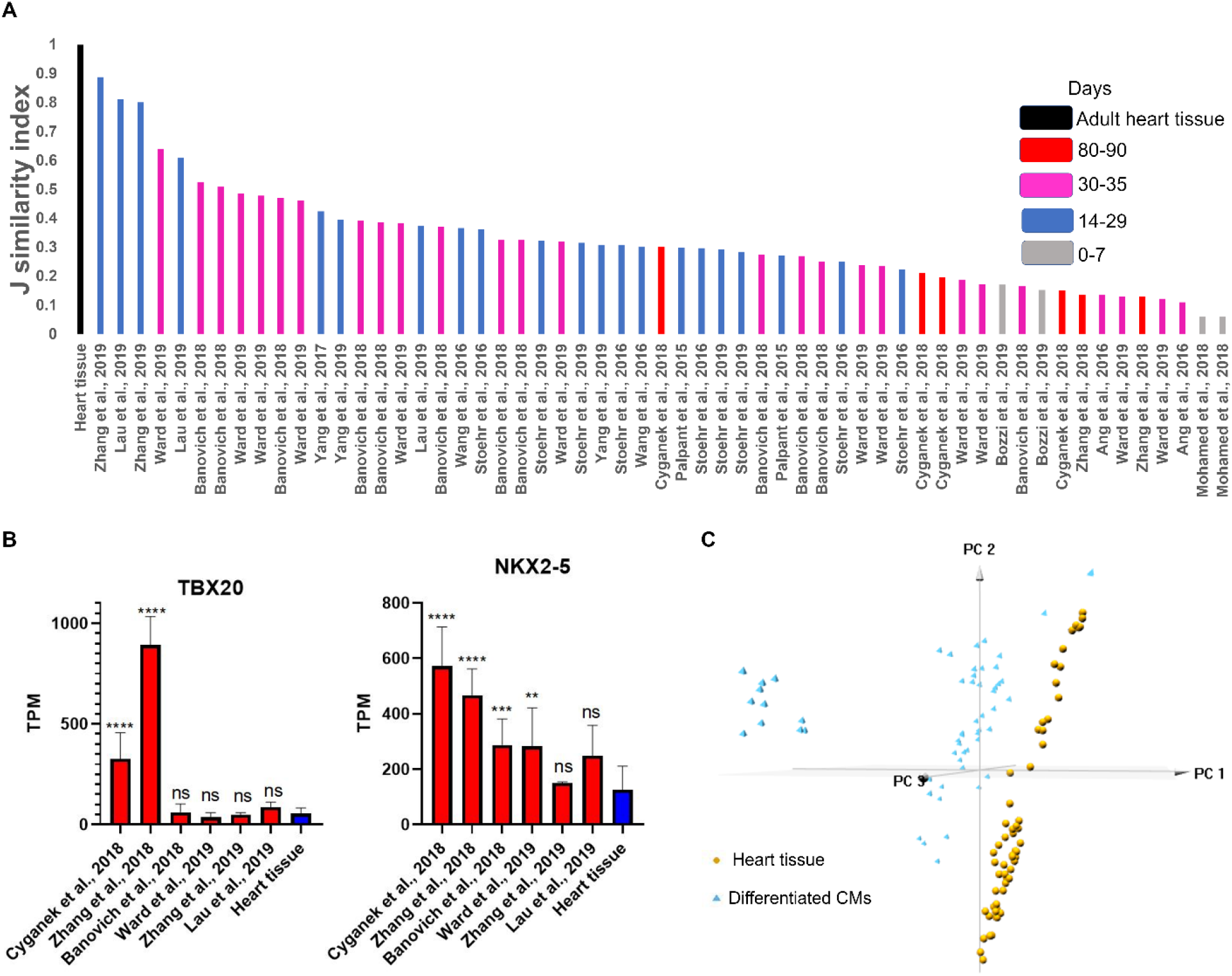
A) Jaccard similarity index which shows resemblance of differentiated CMs to the pooled primary CM samples using expression value of two transcription factors TBX20 and NKX2-5. The similarity value is represented in the Y axis and different differentiation protocols in the X axis. B) TPM values of TBX20 and NKX2-5 expression in six different protocols and in the heart tissue. C) Principle component analysis of the primary heart tissue samples and all differentiated cardiomyocytes (CMs). Primary cardiac tissue and differentiated CMs are represented in blue and yellow respectively.

**Fig S8.**
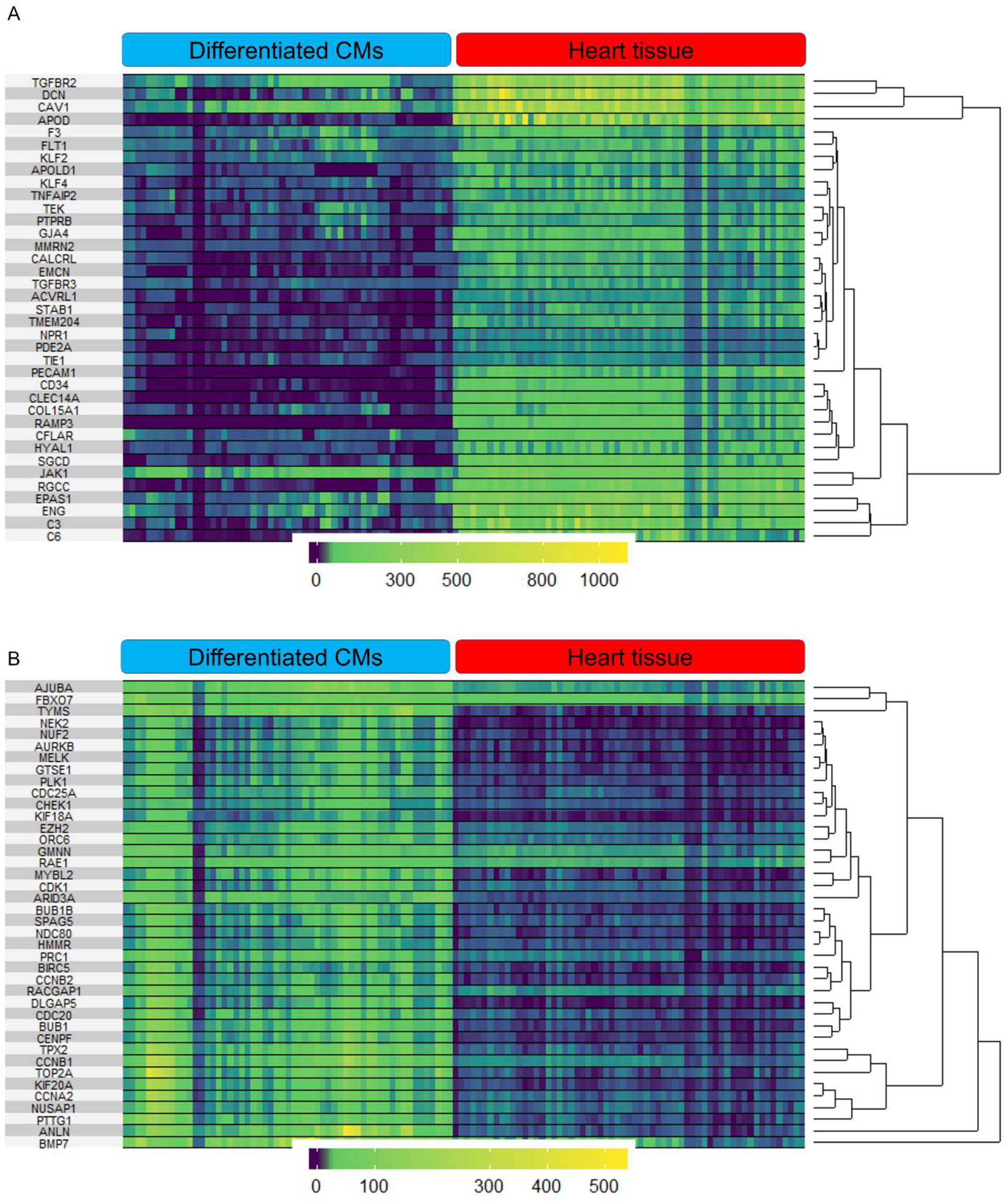
The heatmap plots show expression of 40 genes that have higher A) and lower B) expression in primary CMs compared to the differentiated CM. The yellow and dark blue colors show high and low expression of the genes. The TPM values were used to generate the heatmap using superheat package.

**Fig S9.**
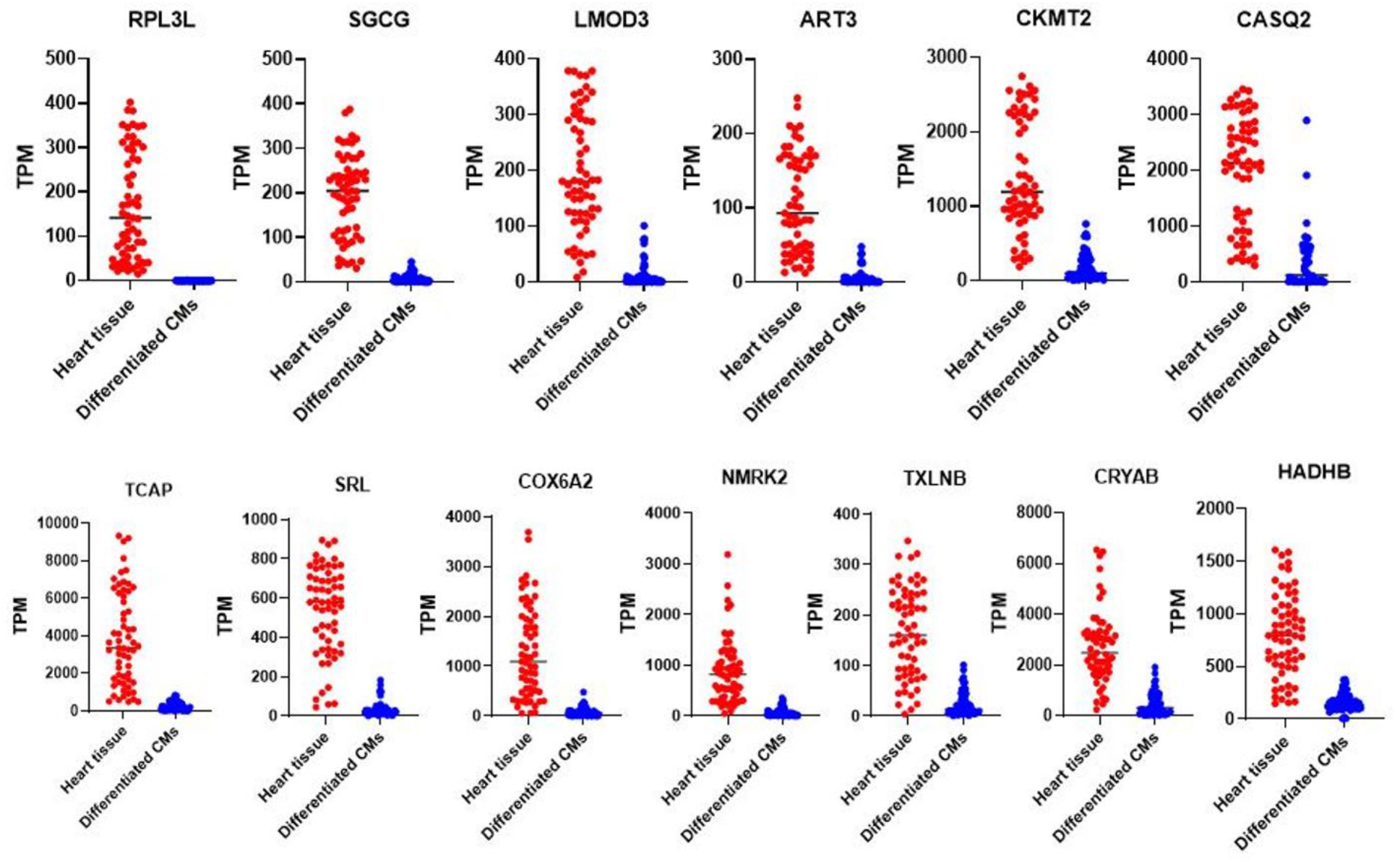
Expression of the genes have higher expression in primary CMs compared to the differentiated ones and are among the list of CM-specific genes (118 genes). The TPM values were used to plot the expression of genes.

**Fig S10.**
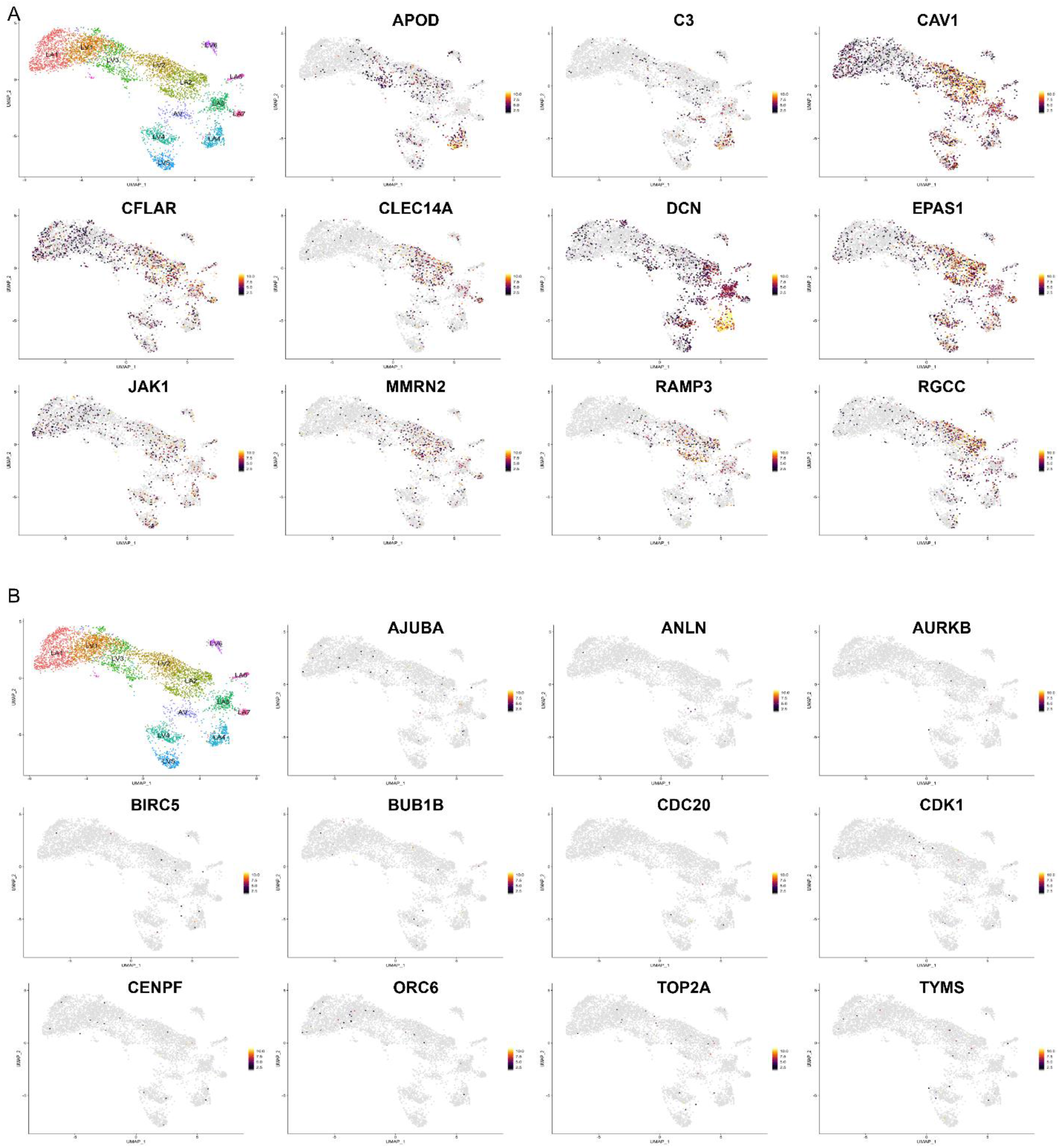
The expression of genes that have higher A) and lower B) expression in primary CMs compared to the differentiated CMs are represented. The expression of genes is indicated by a color gradient in which yellow show high and grey shows low expression.

**Fig S11.**
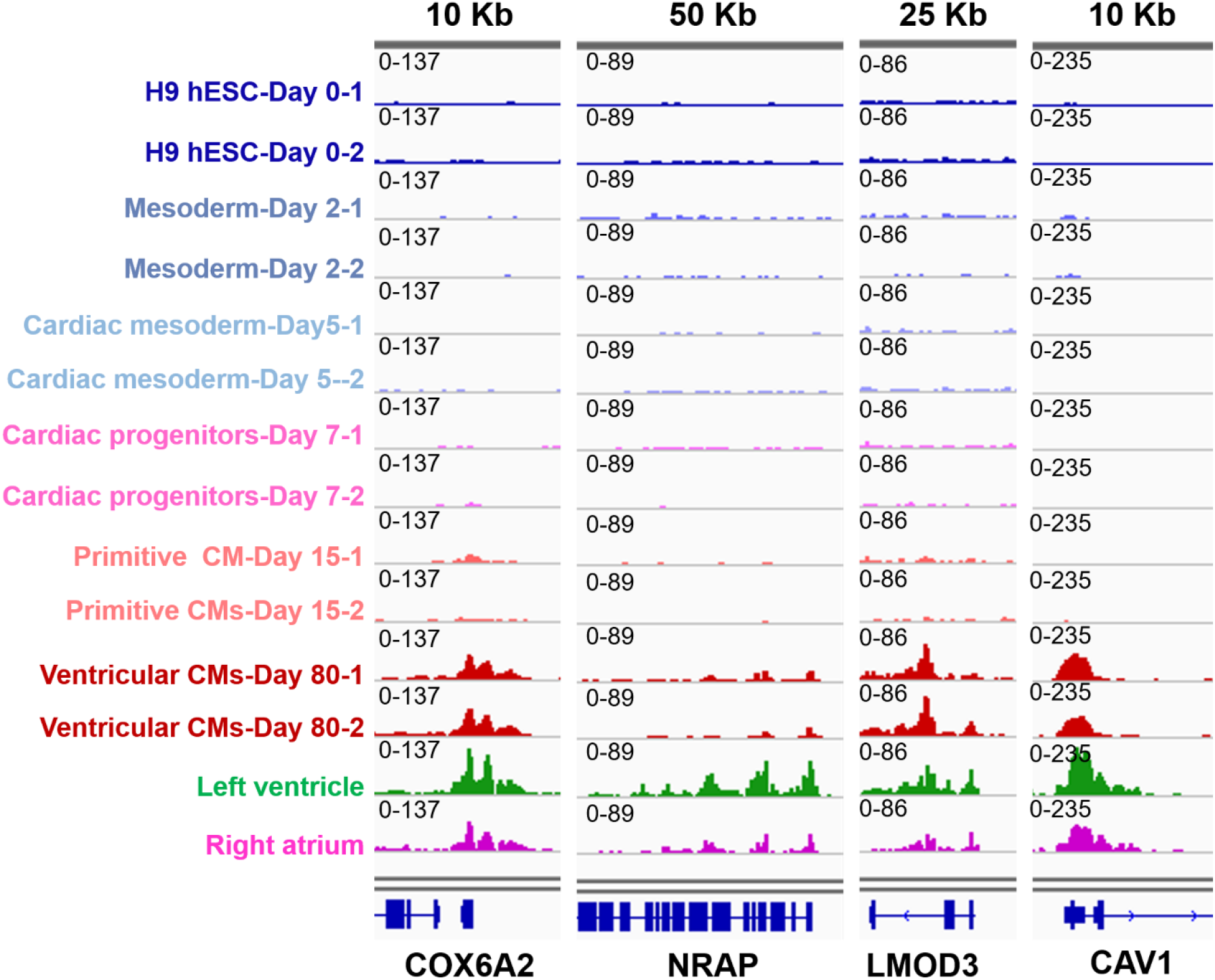
A) H3K27ac peaks on the promotor region of the four CM-specific genes. The length of the covered region is represented above each column B) The schematic procedure of meta-analysis approach for human RNA-seq samples. Somatic cells were compared to the CMs, five cells at a time in the form of 81 comparisons. Statistical parameters were selected to define the list of differentially expressed genes.

**Fig S12.**
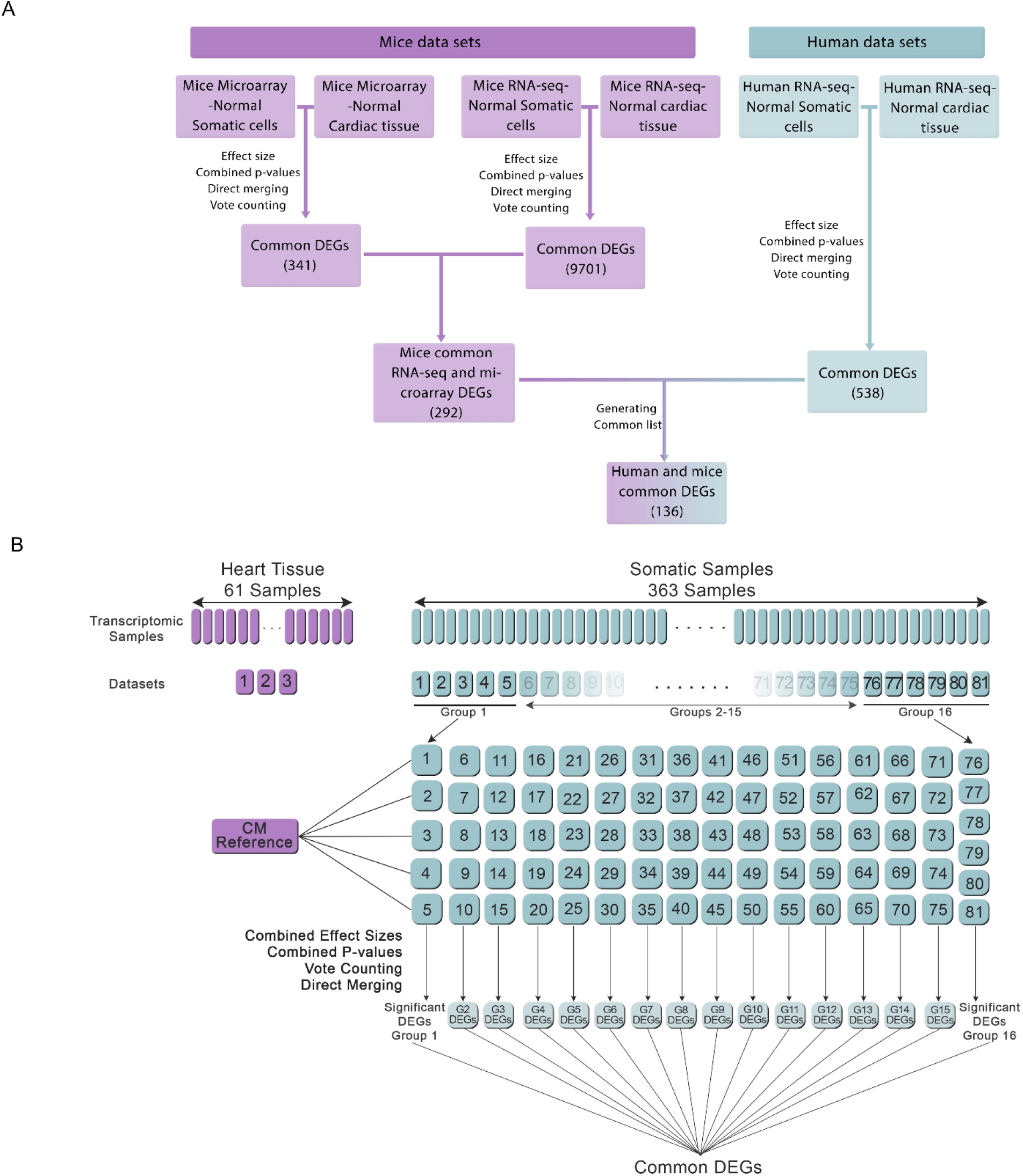
A) Overall representation of the gathered samples, the criteria which were used to detect differentially expressed genes (DEGs) and their number. B) The schematic representation of the experimental design in our study.

## References

1 Peter, A. K., Bjerke, M. A. & Leinwand, L. A. Biology of the cardiac myocyte in heart disease. Mol Biol Cell 27, 2149–2160, doi:10.1091/mbc.E16-01-0038 (2016).

2 Roth, G. A. et al. Global, regional, and national age-sex-specific mortality for 282 causes of death in 195 countries and territories, 1980–2017: a systematic analysis for the Global Burden of Disease Study 2017. The Lancet 392, 1736–1788, doi:10.1016/S0140-6736(18)32203-7 (2018).

3 Kyu, H. H. et al. Global, regional, and national disability-adjusted life-years (DALYs) for 359 diseases and injuries and healthy life expectancy (HALE) for 195 countries and territories, 1990–2017: a systematic analysis for the Global Burden of Disease Study 2017. The Lancet 392, 1859–1922, doi:10.1016/S0140-6736(18)32335-3 (2018).

4 Uhlén, M. et al. Transcriptomics resources of human tissues and organs. Molecular Systems Biology 12, 862, doi:10.15252/msb.20155865 (2016).

5 Wang, L. et al. Single-cell reconstruction of the adult human heart during heart failure and recovery reveals the cellular landscape underlying cardiac function. Nature Cell Biology 22, 108–119, doi:10.1038/s41556-019-0446-7 (2020).

6 DeLaughter, D. M. et al. Single-Cell Resolution of Temporal Gene Expression during Heart Development. Dev Cell 39, 480–490, doi:10.1016/j.devcel.2016.10.001 (2016).

7 Chaudhry, F. et al. Single-Cell RNA Sequencing of the Cardiovascular System: New Looks for Old Diseases. Front Cardiovasc Med 6, 173–173, doi:10.3389/fcvm.2019.00173 (2019).

8 Skelly, D. A. et al. Single-Cell Transcriptional Profiling Reveals Cellular Diversity and Intercommunication in the Mouse Heart. Cell Reports 22, 600–610, doi:10.1016/j.celrep.2017.12.072. (2018).

9 Alba, A. et al. Complications after heart transplantation: hope for the best, but prepare for the worst. Int J Transplant Res Med 2, 2–22 (2016).

10 Später, D., Hansson, E. M., Zangi, L. & Chien, K. R. How to make a cardiomyocyte. Development 141, 4418, doi:10.1242/dev.091538 (2014).

11 Lee Richard, T. & Walsh, K. The Future of Cardiovascular Regenerative Medicine. Circulation 133, 2618–2625, doi:10.1161/CIRCULATIONAHA.115.019214 (2016).

12 Witman, N. & Sahara, M. Cardiac Progenitor Cells in Basic Biology and Regenerative Medicine. Stem Cells Int 2018, 8283648–8283648, doi:10.1155/2018/8283648 (2018).

13 van den Berg, C. W., Elliott, D. A., Braam, S. R., Mummery, C. L. & Davis, R. P. Differentiation of Human Pluripotent Stem Cells to Cardiomyocytes Under Defined Conditions. In: Nagy A., Turksen K. (eds) Patient-Specific Induced Pluripotent Stem Cell Models. Methods in Molecular Biology 1353. Humana Press, New York, NY (2014).

14 Ieda, M. et al. Direct Reprogramming of Fibroblasts into Functional Cardiomyocytes by Defined Factors. Cell 142, 375–386, doi:10.1016/j.cell.2010.07.002 (2010).

15 Kolanowski, T. J., Antos, C. L. & Guan, K. Making human cardiomyocytes up to date: Derivation, maturation state and perspectives. International Journal of Cardiology 241, 379–386, doi:10.1016/j.ijcard.2017.03.099 (2017).

16 van den Berg, C. W. et al. Transcriptome of human foetal heart compared with cardiomyocytes from pluripotent stem cells. Development 142, 3231–3238, doi:10.1242/dev.123810 (2015).

17 Veerman, C. C. et al. Immaturity of Human Stem-Cell-Derived Cardiomyocytes in Culture: Fatal Flaw or Soluble Problem? Stem Cells and Development 24, 1035–1052, doi:10.1089/scd.2014.0533 (2015).

18 van den Heuvel, N. H. L., van Veen, T. A. B., Lim, B. & Jonsson, M. K. B. Lessons from the heart: Mirroring electrophysiological characteristics during cardiac development to in vitro differentiation of stem cell derived cardiomyocytes. Journal of Molecular and Cellular Cardiology 67, 12–25, doi:10.1016/j.yjmcc.2013.12.011 (2014).

19 Waas, M. et al. Are These Cardiomyocytes? Protocol Development Reveals Impact of Sample Preparation on the Accuracy of Identifying Cardiomyocytes by Flow Cytometry. Stem Cell Reports 12, 395–410, doi:10.1016/j.stemcr.2018.12.016 (2019).

20 Schaum, N. et al. Single-cell transcriptomics of 20 mouse organs creates a Tabula Muris. Nature 562, 367–372, doi:10.1038/s41586-018-0590-4 (2018).

21 Uhlén, M. et al. Tissue-based map of the human proteome. Science 347, 1260419, doi:10.1126/science.1260419 (2015).

22 Kundaje, A. et al. Integrative analysis of 111 reference human epigenomes. Nature 518, 317–330, doi:10.1038/nature14248 (2015).

23 Yoshida, Y. & Yamanaka, S. Induced Pluripotent Stem Cells 10 Years Later. Circulation Research 120, 1958–1968, doi:10.1161/CIRCRESAHA.117.311080 (2017).

24 Lian, X. et al. Directed cardiomyocyte differentiation from human pluripotent stem cells by modulating Wnt/β-catenin signaling under fully defined conditions. Nat Protoc 8, 162–175, doi:10.1038/nprot.2012.150 (2012).

25 Burridge, P. W. et al. Chemically defined generation of human cardiomyocytes. Nat Methods 11, 855–860, doi:10.1038/nmeth.2999 (2014).

26 Elliott, D. A. et al. NKX2-5eGFP/w hESCs for isolation of human cardiac progenitors and cardiomyocytes. Nat Methods 8, 1037–1040, doi:10.1038/nmeth.1740 (2011).

27 Zhang, J. et al. Functional cardiac fibroblasts derived from human pluripotent stem cells via second heart field progenitors. Nature Communications 10, doi:10.1038/s41467-019-09831-5 (2019).

28 Banovich, N. E. et al. Impact of regulatory variation across human iPSCs and differentiated cells. Genome Res 28, 122–131, doi:10.1101/gr.224436.117 (2018).

29 Ang, Y.-S. et al. Disease Model of GATA4 Mutation Reveals Transcription Factor Cooperativity in Human Cardiogenesis. Cell 167, 1734–1749.e1722, doi:10.1016/j.cell.2016.11.033 (2016).

30 Palpant, N. J. et al. Inhibition of β-catenin signaling respecifies anterior-like endothelium into beating human cardiomyocytes. Development 142, 3198, doi:10.1242/dev.117010 (2015).

31 Cyganek, L. et al. Deep phenotyping of human induced pluripotent stem cell-derived atrial and ventricular cardiomyocytes. JCI Insight 3, e99941, doi:10.1172/jci.insight.99941 (2018).

32 Yang, X. et al. Fatty Acids Enhance the Maturation of Cardiomyocytes Derived from Human Pluripotent Stem Cells. Stem Cell Reports 13, 657–668, doi:10.1016/j.stemcr.2019.08.013 (2019).

33 Kattman, S. J. et al. Stage-Specific Optimization of Activin/Nodal and BMP Signaling Promotes Cardiac Differentiation of Mouse and Human Pluripotent Stem Cell Lines. Cell Stem Cell 8, 228–240, doi:10.1016/j.stem.2010.12.008 (2011).

34 Bertero, A. et al. Dynamics of genome reorganization during human cardiogenesis reveal an RBM20-dependent splicing factory. Nature Communications 10, 1538, doi:10.1038/s41467-019-09483-5 (2019).

35 Zhang, Y. et al. Transcriptionally active HERV-H retrotransposons demarcate topologically associating domains in human pluripotent stem cells. Nature Genetics 51, 1380–1388, doi:10.1038/s41588-019-0479-7 (2019).

36 Ward, M. C. & Gilad, Y. A generally conserved response to hypoxia in iPSC-derived cardiomyocytes from humans and chimpanzees. Elife 8, e42374, doi:10.7554/eLife.42374 (2019).

37 Bozzi, A. et al. Using Human Induced Pluripotent Stem Cell-Derived Cardiomyocytes as a Model to Study Trypanosoma cruzi Infection. Stem Cell Reports 12, 1232–1241, doi:10.1016/j.stemcr.2019.04.017 (2019).

38 Stoehr, A. et al. The ribosomal prolyl-hydroxylase OGFOD1 decreases during cardiac differentiation and modulates translation and splicing. JCI Insight 5, e128496, doi:10.1172/jci.insight.128496 (2019).

39 Wang, X. et al. Discovery and validation of sub-threshold genome-wide association study loci using epigenomic signatures. Elife 5, e10557, doi:10.7554/eLife.10557 (2016).

40 Stoehr, A. et al. Prolyl hydroxylation regulates protein degradation, synthesis, and splicing in human induced pluripotent stem cell-derived cardiomyocytes. Cardiovascular Research 110, 346–358, doi:10.1093/cvr/cvw081 (2016).

41 Yang, C. et al. Induced pluripotent stem cell modelling of HLHS underlines the contribution of dysfunctional NOTCH signalling to impaired cardiogenesis. Human Molecular Genetics 26, 3031–3045, doi:10.1093/hmg/ddx140 (2017).

42 Mohamed, T. M. A. et al. Regulation of Cell Cycle to Stimulate Adult Cardiomyocyte Proliferation and Cardiac Regeneration. Cell 173, 104–116.e112, doi:10.1016/j.cell.2018.02.014 (2018).

43 Lau, E. et al. Splice-Junction-Based Mapping of Alternative Isoforms in the Human Proteome. Cell Reports 29, 3751–3765.e3755, doi:10.1016/j.celrep.2019.11.026 (2019).

44 Greulich, F., Rudat, C. & Kispert, A. Mechanisms of T-box gene function in the developing heart. Cardiovascular Research 91, 212–222, doi:10.1093/cvr/cvr112 (2011).

45 Kanatous, S. B. & Mammen, P. P. A. Regulation of myoglobin expression. The Journal of Experimental Biology 213, 2741, doi:10.1242/jeb.041442 (2010).

46 Hüttemann, M. et al. Mice deleted for heart-type cytochrome c oxidase subunit 7a1 develop dilated cardiomyopathy. Mitochondrion 12, 294–304, doi:10.1016/j.mito.2011.11.002 (2012).

47 Truszkowska, G. T. et al. Homozygous truncating mutation in NRAP gene identified by whole exome sequencing in a patient with dilated cardiomyopathy. Sci Rep 7, doi:10.1038/s41598-017-03189-8 (2017).

48 Sun, Q. et al. SEMA6D regulates perinatal cardiomyocyte proliferation and maturation in mice. Developmental Biology 452, 1–7, doi:10.1016/j.ydbio.2019.04.013 (2019).

49 Danielsson, F., James, T., Gomez-Cabrero, D. & Huss, M. Assessing the consistency of public human tissue RNA-seq data sets. Briefings in Bioinformatics 16, 941–949, doi:10.1093/bib/bbv017 (2015).

50 Arora, S., Pattwell, S. S., Holland, E. C. & Bolouri, H. Variability in estimated gene expression among commonly used RNA-seq pipelines. Scientific Reports 10, 2734, doi:10.1038/s41598-020-59516-z (2020).

51 Lonsdale, J. et al. The Genotype-Tissue Expression (GTEx) project. Nature Genetics 45, 580–585, doi:10.1038/ng.2653 (2013).

52 Anderson, D. J. et al. NKX2-5 regulates human cardiomyogenesis via a HEY2 dependent transcriptional network. Nature Communications 9, doi:10.1038/s41467-018-03714-x (2018).

53 Singh, M. K. et al. Tbx20 is essential for cardiac chamber differentiation and repression of Tbx2. Development (Cambridge, England) 132, 2697–2707, doi:10.1242/dev.01854 (2005).

54 Wiegering, A., Rüther, U. & Gerhardt, C. The Role of Hedgehog Signalling in the Formation of the Ventricular Septum. J Dev Biol 5, doi:10.3390/jdb5040017 (2017).

55 Man, J., Barnett, P. & Christoffels, V. M. Structure and function of the Nppa-Nppb cluster locus during heart development and disease. Cell Mol Life Sci 75, 1435–1444, doi:10.1007/s00018-017-2737-0 (2018).

56 Cheng, C. et al. Mutation in NPPA causes atrial fibrillation by activating inflammation and cardiac fibrosis in a knock-in rat model. The FASEB Journal 33, 8878–8891, doi:10.1096/fj.201802455RRR (2019).

57 Warren, S. A. et al. Differential Role of Nkx2-5 in Activation of the Atrial Natriuretic Factor Gene in the Developing versus Failing Heart. Molecular and Cellular Biology 31, 4633–4645, doi:10.1128/MCB.05940-11 (2011).

58 Brown, D. D. et al. Tbx5 and Tbx20 act synergistically to control vertebrate heart morphogenesis. Development (Cambridge, England) 132, 553–563, doi:10.1242/dev.01596 (2005).

59 Carroll, S. L. & Horowits, R. Myofibrillogenesis and formation of cell contacts mediate the localization of N-RAP in cultured chick cardiomyocytes. Cell Motility 47, 63–76, doi:10.1002/1097-0169(200009)47:1<63::AID-CM6>3.0.CO;2-M (2000).

60 Mortazavi, A., Williams, B. A., McCue, K., Schaeffer, L. & Wold, B. Mapping and quantifying mammalian transcriptomes by RNA-Seq. Nature Methods 5, 621–628, doi:10.1038/nmeth.1226 (2008).

61 Wagner, G. P., Kin, K. & Lynch, V. J. Measurement of mRNA abundance using RNA-seq data: RPKM measure is inconsistent among samples. Theory in Biosciences 131, 281–285, doi:10.1007/s12064-012-0162-3 (2012).

62 Wettenhall, J. M., Simpson, K. M., Satterley, K. & Smyth, G. K. affylmGUI: a graphical user interface for linear modeling of single channel microarray data. Bioinformatics 22, 897–899, doi:10.1093/bioinformatics/btl025 (2006).

63 Irizarry, R. A. et al. Exploration, normalization, and summaries of high density oligonucleotide array probe level data. Biostatistics 4, 249–264, doi:10.1093/biostatistics/4.2.249 (2003).

64 Robinson, J. T. et al. Integrative genomics viewer. Nature Biotechnology 29, 24–26, doi:10.1038/nbt.1754 (2011).

65 Thorvaldsdóttir, H., Robinson, J. T. & Mesirov, J. P. Integrative Genomics Viewer (IGV): high-performance genomics data visualization and exploration. Brief Bioinform 14, 178–192, doi:10.1093/bib/bbs017 (2013).

66 Zhou, G. et al. NetworkAnalyst 3.0: a visual analytics platform for comprehensive gene expression profiling and meta-analysis. Nucleic Acids Research 47, W234–W241, doi:10.1093/nar/gkz240 (2019).

67 Butler, A., Hoffman, P., Smibert, P., Papalexi, E. & Satija, R. Integrating single-cell transcriptomic data across different conditions, technologies, and species. Nature Biotechnology 36, 411–420, doi:10.1038/nbt.4096 (2018).

68 Stuart, T. et al. Comprehensive Integration of Single-Cell Data. Cell 177, 1888–1902.e1821, doi:10.1016/j.cell.2019.05.031 (2019).

69 Chen, E. Y. et al. Enrichr: interactive and collaborative HTML5 gene list enrichment analysis tool. BMC Bioinformatics 14, 128, doi:10.1186/1471-2105-14-128 (2013).

70 Kuleshov, M. V. et al. Enrichr: a comprehensive gene set enrichment analysis web server 2016 update. Nucleic Acids Research 44, W90–W97, doi:10.1093/nar/gkw377 (2016).

71 Scardoni, G., Petterlini, M. & Laudanna, C. Analyzing biological network parameters with CentiScaPe. Bioinformatics (Oxford, England) 25, 2857–2859, doi:10.1093/bioinformatics/btp517 (2009).

72 Shannon, P. et al. Cytoscape: a software environment for integrated models of biomolecular interaction networks. Genome Res 13, 2498–2504, doi:10.1101/gr.1239303 (2003).

73 Szklarczyk, D. et al. STRING v11: protein–protein association networks with increased coverage, supporting functional discovery in genome-wide experimental datasets. Nucleic Acids Research 47, D607–D613, doi:10.1093/nar/gky1131 (2018).

74 Saito, R. et al. A travel guide to Cytoscape plugins. Nat Methods 9, 1069–1076, doi:10.1038/nmeth.2212 (2012).

75 Bader, G. D. & Hogue, C. W. V. An automated method for finding molecular complexes in large protein interaction networks. BMC bioinformatics 4, doi:10.1186/1471-2105-4-2 (2003).

76 Raudvere, U. et al. g:Profiler: a web server for functional enrichment analysis and conversions of gene lists (2019 update). Nucleic Acids Research 47, W191–W198, doi:10.1093/nar/gkz369 (2019).

77 Barter, R. L. & Yu, B. Superheat: An R package for creating beautiful and extendable heatmaps for visualizing complex data. J Comput Graph Stat 27, 910–922, doi:10.1080/10618600.2018.1473780 (2018).

